# Type I and Type III Interferons Differentially Shape Antiviral Defense and Epithelial Integrity at the Choroid Plexus

**DOI:** 10.64898/2026.02.10.705109

**Authors:** Justin T. Steppe, Emma Heckenberg, Caitlin Hale, Carolyn B. Coyne

## Abstract

The choroid plexus (ChP) forms the blood-cerebrospinal fluid (CSF) barrier, yet how this CNS interface responds to viral infection remains poorly understood. Here, we identify the ChP epithelium as a major target of echovirus infection and establish complementary human organoid and mouse models to define antiviral responses at this barrier. Single-cell RNA sequencing of cerebral organoids revealed preferential infection of ChP epithelial cells, and ChP organoids supported productive infection accompanied by a robust type III interferon (IFN) response. In mice expressing human FcRn, the echovirus receptor, infection similarly targeted the ChP, enabling genetic dissection of IFN signaling *in vivo*. Type I IFN signaling restricted viral replication and protected against lethal disease. In contrast, type III IFN signaling was dispensable for viral control but promoted ChP epithelial injury and, following viral clearance, was associated with persistent motor deficits, ependymal loss, and ventriculomegaly. These findings reveal distinct functions for type I and III IFNs at the blood-CSF barrier and identify type III IFN signaling as a driver of tissue injury and lasting neurological sequelae following viral infection.

## Introduction

The choroid plexus (ChP) is a specialized secretory tissue located in each ventricle of the brain. It consists of a highly folded layer of simple cuboidal epithelial cells surrounding a stroma of fenestrated capillaries and connective tissue. The ChP performs two essential functions: the production of cerebrospinal fluid (CSF) and the formation of the blood-CSF barrier, which protects the brain against systemic insults. The ChP epithelial barrier is established by tight junctions, which selectively regulate the permeability of molecules from the circulation into the CSF ^1^. Along with coordinated secretion of ChP-derived proteins, this control of permeability is critical in maintaining homeostasis of the CSF ^2^. The ChP also plays a major role in coordinating the trafficking of immune cells into the brain in response to infection or injury ^3^. Despite its crucial role in controlling access to the brain, there is limited knowledge about how the ChP responds to infections, particularly by viruses.

Viral infections of the central nervous system (CNS), even when clinically resolved, can lead to lasting neurological sequelae across the lifespan, including neurodevelopmental deficits, structural abnormalities, and long-term cognitive impairment ^4-6^. Although the routes by which viruses access the CNS remain incompletely defined, the ChP has been implicated as a potential portal of entry for several pathogens, including HIV, SARS-CoV-2, Zika virus, and some enteroviruses ^7-11^. For some viruses, interaction with the ChP occurs without direct infection of the epithelium, whereas others may actively replicate within the epithelium ^7-11^. ChP epithelial cells can mount robust immune responses to pathogen associated molecular patterns (PAMPs) such as LPS, yet their responses to active viral infection remain poorly characterized ^12^. This gap in knowledge reflects, in part, the historical lack of tractable experimental systems to model viral infection at the blood-CSF interface. Recently, ChP organoids have emerged as a powerful *in vitro* platform to study host pathogen interactions at this critical barrier site ^13^.

Enteroviruses are a leading cause of aseptic meningitis worldwide, with echoviruses among the most frequently detected etiologic agents ^14,15^. Echovirus RNA is readily detected in the CSF of patients with meningitis, indicating that these viruses can access the CSF compartment during infection ^15^. The ChP epithelium expresses high levels of the neonatal Fc receptor (FcRn), the primary entry receptor for echoviruses, positioning this tissue as a plausible site of viral entry into and interaction with the CSF compartment ^16-18^. Consistent with this possibility, echovirus 30 can directly infect ChP epithelial cells *in vitro* ^11^. However, how the ChP responds to echovirus infection *in vivo*, which antiviral pathways control infection at this barrier, and whether these responses protect or injure the ChP remain unknown. Thus, echoviruses provide a clinically relevant system in which to define antiviral immunity at the blood-CSF barrier and determine how host defense at this interface shapes both viral control and neurological injury.

One of the first host cell defenses against viral infections is the production of interferons (IFNs), including type I and type III IFNs. Both classes of IFNs are potent antiviral cytokines that activate largely overlapping signaling pathways yet differ markedly in receptor distribution. The type I IFN receptor is expressed on nearly all nucleated cells, whereas expression of the type III IFN receptor is largely restricted to epithelial cells and select immune populations ^19^. Type I IFNs play a well-established role in restricting viral infections in the CNS, but far less is known about the functions of type III IFNs in the brain ^20-22^. In peripheral tissues, type III IFNs can exert protective effects by limiting viral replication in barrier epithelia such as the GI tract, yet they can also have detrimental consequences by impairing epithelial repair in the lung and gut ^23-25^. In the nervous system, type III IFNs have been shown to limit neuroinvasion by West Nile virus through strengthening the BBB, but their roles at other CNS interfaces remain largely unexplored ^26^. It is unclear how type I and type III IFN signaling shape antiviral defense and tissue integrity at the ChP during viral infection.

Here, we established complementary *in vitro* and *in vivo* platforms to interrogate viral infection at the ChP using echoviruses as a tractable and clinically relevant model system. Using human ChP organoids, we show that ChP-derived epithelial cells mount a robust type III IFN response to echovirus infection. In parallel, by developing a mouse model of echovirus infection that targets the ChP *in vivo*, we uncover a striking functional divergence between IFN pathways at this CNS interface. While type I IFNs are indispensable for controlling viral replication, type III IFNs impair barrier repair, exacerbate epithelial injury, and worsen sequelae following infection. Together, these findings redefine IFN function at the blood-CSF interface by uncovering opposing roles for type I and type III IFNs in antiviral defense and tissue repair.

## Results

### Echovirus Preferentially Infects the Choroid Plexus Epithelium in Human Cerebral Organoids

To determine whether echoviruses exhibit selective tropism for the ChP epithelium in an unbiased human system, we generated cerebral organoids using a previously established unguided differentiation protocol and infected them with echovirus 5 (E5) ^27^. We focused on day 70 organoids, which reproducibly develop a well-defined ChP epithelial compartment alongside diverse neuronal and progenitor populations. Immunohistochemical (IHC) analysis confirmed the presence of mature neural lineages and progenitor cells, as indicated by expression of MAP2 and SOX2, respectively (**Figure 1A** and **1B**), and immunofluorescence staining further revealed organized ChP epithelium marked by transthyretin (TTR), adjacent to surrounding neuronal tissue marked by βIII-tubulin (TUBB3) (**Figure 1C**). Together, these data establish that these organoids contain the cellular architecture necessary to model viral infection at the ChP within a complex neural environment, providing a physiologically relevant context to assess echovirus tropism across multiple lineages.

**Figure 1.**
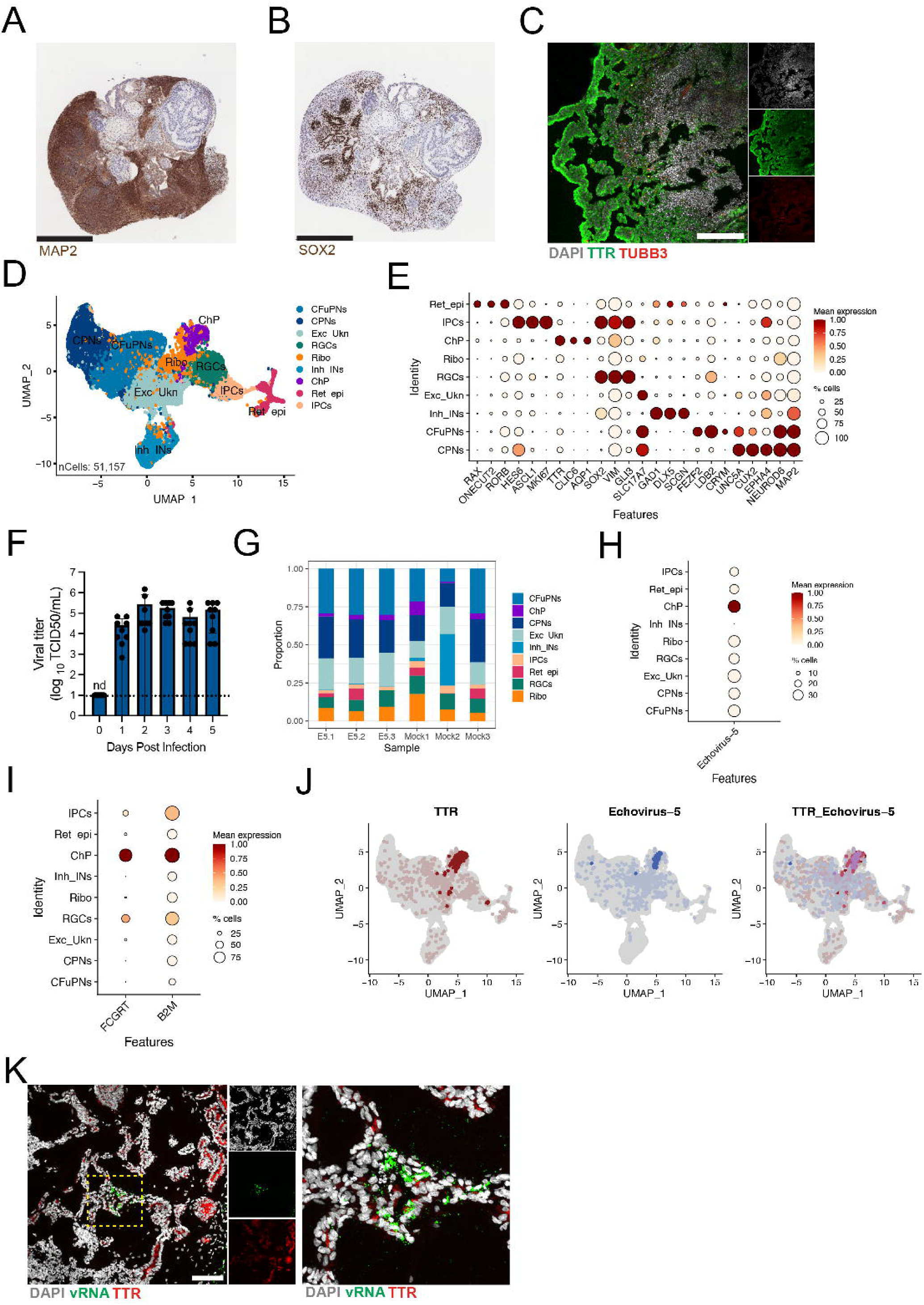
Echovirus tropism in human cerebral organoids. All cerebral organoids used were 70±5 days old **(A-K)**. **(A)** Immunohistochemical staining of cerebral organoids for MAP2. **(B)** Immunohistochemical staining for SOX2. **(C)** Immunofluorescence staining for DAPI (gray), transthyretin (TTR; green), and beta tubulin III (TUBB3; red). **(D)** UMAP of scRNA-seq from three E5-infected and three mock-infected cerebral organoids collected at four days post infection, with annotated cell types including callosal projection neurons (CPNs), corticofugal projection neurons (CFuPNs), inhibitory interneurons (Inh_INs), excitatory neurons of unknown identity (Exc_Ukn), radial glial cells (RGCs), ribosome-high cells (Ribo), choroid plexus epithelium (ChP), intermediate progenitor cells (IPCs), and retinal epithelial cells (Ret_epi). **(E)** Dot plot showing expression of selected marker genes for each annotated cell type. **(F)** Viral titers in supernatants from cerebral organoids infected with 200 PFU of E5 virus over a five-day time course, measured by TCID50 assay and shown as log_10_ TCID_50_ per milliliter; the dotted line indicates the limit of detection, with data shown as mean ± standard deviation and individual organoids represented as points. **(G)** Proportion of cells assigned to each cluster, shown by individual sample. **(H)** Dot plot showing detection of E5 viral transcripts across cell clusters. **(I)** Dot plot showing expression of the neonatal Fc receptor (FCGRT) and β-2 microglobulin (β2M) across clusters. **(J)** Merged feature plot showing expression of TTR (red) and E5 viral transcripts (blue); purple indicates cells expressing both features. **(K)** Immunofluorescence staining of cerebral organoid section at three days post infection for DAPI (gray), viral RNA (green), and TTR (red). Scale bars are 500 µm (A, B), 200 µm (C), and 100 µm (K). For dot plots, values are scaled such that 1 represents the highest normalized expression and 0 represents the lowest normalized expression, and dot size indicates the percentage of cells within each cluster expressing the indicated gene.

To more comprehensively define the cellular composition of these organoids and identify cell types permissive to infection, we performed single cell RNA sequencing (scRNA-seq) on three mock infected and three E5 infected organoids. Organoids were manually dissociated and processed in parallel to minimize technical variability. Sequencing reads were aligned to a custom reference genome containing both the human genome (GRCh38) and the E5 genome. To account for inter-sample variation, datasets were integrated using Harmony batch correction ^28^, and the number of detected features, total UMI counts, and percentage of mitochondrial transcripts were regressed during normalization. After quality control filtering, a total of 51,157 cells were retained and visualized using UMAP projections (**Figure 1D**). Unsupervised clustering identified nine distinct cell populations based on canonical marker gene expression (**Figure 1D, 1E**, **Supplemental Table 1**). These included multiple immature neuronal populations, such as callosal projection neurons expressing *UNC5A*, *CUX2*, and *EPHA4*, corticofugal projection neurons expressing *FEZF2*, *LDB2*, and *CRYM*, inhibitory interneurons expressing *GAD1*, *DLX5*, and *SCGN*, and an excitatory neuronal population marked by *SLC17A7* but lacking other distinguishing markers. Callosal projection neurons form excitatory connections between the cortical hemispheres, while corticofugal projection neurons form excitatory connections between the cortex and subcortical structures, and inhibitory interneurons form critical inhibitory connections with these excitatory projection neurons. Neural progenitor populations were also identified, including radial glial cells expressing *SOX2*, *VIM*, and *GLI3* and intermediate progenitor cells expressing *HES6*, *ASCL1*, and *MKI67*. In addition, we detected several non-neuronal populations, including ChP epithelial cells marked by *TTR*, *AQP1*, and *CLIC6*, retinal epithelial cells expressing *RAX*, *ONECUT2*, and *RORB*, and a cluster characterized by elevated expression of ribosomal genes. Together, these data confirm that cerebral organoids recapitulate a broad spectrum of cell types present in the developing human brain, enabling an unbiased assessment of E5 tropism across neural and barrier associated lineages.

Having defined the cellular composition of cerebral organoids by scRNA-seq, we next asked whether these organoids were permissive to echovirus infection. To address this, cerebral organoids were infected with 200 PFU of E5 virus and monitored over a five-day time course. Experiments were performed using three independent organoid batches, with three organoids per condition per batch, to account for batch-to-batch variability. High titers of infectious virus were detected in the supernatant throughout the time course, indicating productive infection (**Figure 1F**). We then assessed whether echovirus infection altered overall cellular composition. Cluster proportions across samples showed no major differences between mock- and E5-infected organoids (**Figure 1G**). Some variability was observed between individual organoids, with inhibitory interneurons and retinal epithelial cells not consistently detected across all samples (**Figure 1G**, **Supplemental Figure 1A-B**), consistent with expected heterogeneity in unguided differentiation protocols ^29^.

Finally, we assessed the distribution of viral genome reads across cell populations in cerebral organoids and found that viral transcripts mapped almost exclusively to the ChP epithelial cell cluster (**Figure 1H**). Because receptor expression is a key determinant of cellular susceptibility to viral infection, we examined expression of the echovirus entry receptor, the neonatal Fc receptor (FcRn), encoded by *FCGRT*. We also analyzed expression of β-2 microglobulin (*β2M)*, which is required for FcRn cell surface localization ^30^. The ChP epithelial cells, intermediate progenitor cells, and the radial glial cells all expressed *FCGRT* and *B2M*, however the ChP epithelial cell cluster had the highest expression of both genes among these populations (**Figure 1I**). Consistent with this, merged feature plots demonstrated that viral transcripts localized predominantly to choroid plexus epithelial cells, as indicated by their co-expression with the ChP marker *TTR* (**Figure 1J**). Finally, immunofluorescence staining for double stranded RNA, a marker of active viral replication, together with TTR confirmed productive echovirus infection of the ChP epithelium at three days post infection (**Figure 1K**). We performed additional staining for viral RNA and the neural progenitor cell marker SOX2 but observed very low levels of infection in SOX2^+^ progenitor cells (**Supplemental Figure 1C-D**), consistent with what we observed in the scRNAseq data set. Together, these data demonstrate that in a complex human cerebral organoid model containing diverse neural and non-neural cell types, echoviruses display strong tropism for the ChP epithelium.

### Modeling Choroid Plexus Infection *In Vitro* and *In Vivo* Using Echoviruses

To more directly examine the effects of viral infection on the ChP epithelium, we used a human ChP organoid model ^13^. These organoids recapitulate key features of ChP biology, including expression of canonical epithelial markers, production of CSF, and development of functional barrier properties ^13^. Unless otherwise indicated, day 55 organoids were used for all experiments to ensure appropriate maturation and representation of relevant cell types. We first characterized the structural and molecular features of ChP organoids. In contrast to densely cellular cerebral organoids, ChP organoids formed large cystic structures containing CSF-like fluid (**Figure 2A**). Histological analysis revealed epithelial cells lining these cystic compartments, consistent with ChP architecture (**Figure 2B**). RNAscope and immunohistochemical staining further demonstrated expression of established ChP markers, including TTR and aquaporin-1 (AQP1), as well as the tight junction protein ZO-1 (TJP1) (**Figure 2C**–**E**). Together, these data confirm that the organoids recapitulate the cellular identity of the ChP epithelium.

**Figure 2.**
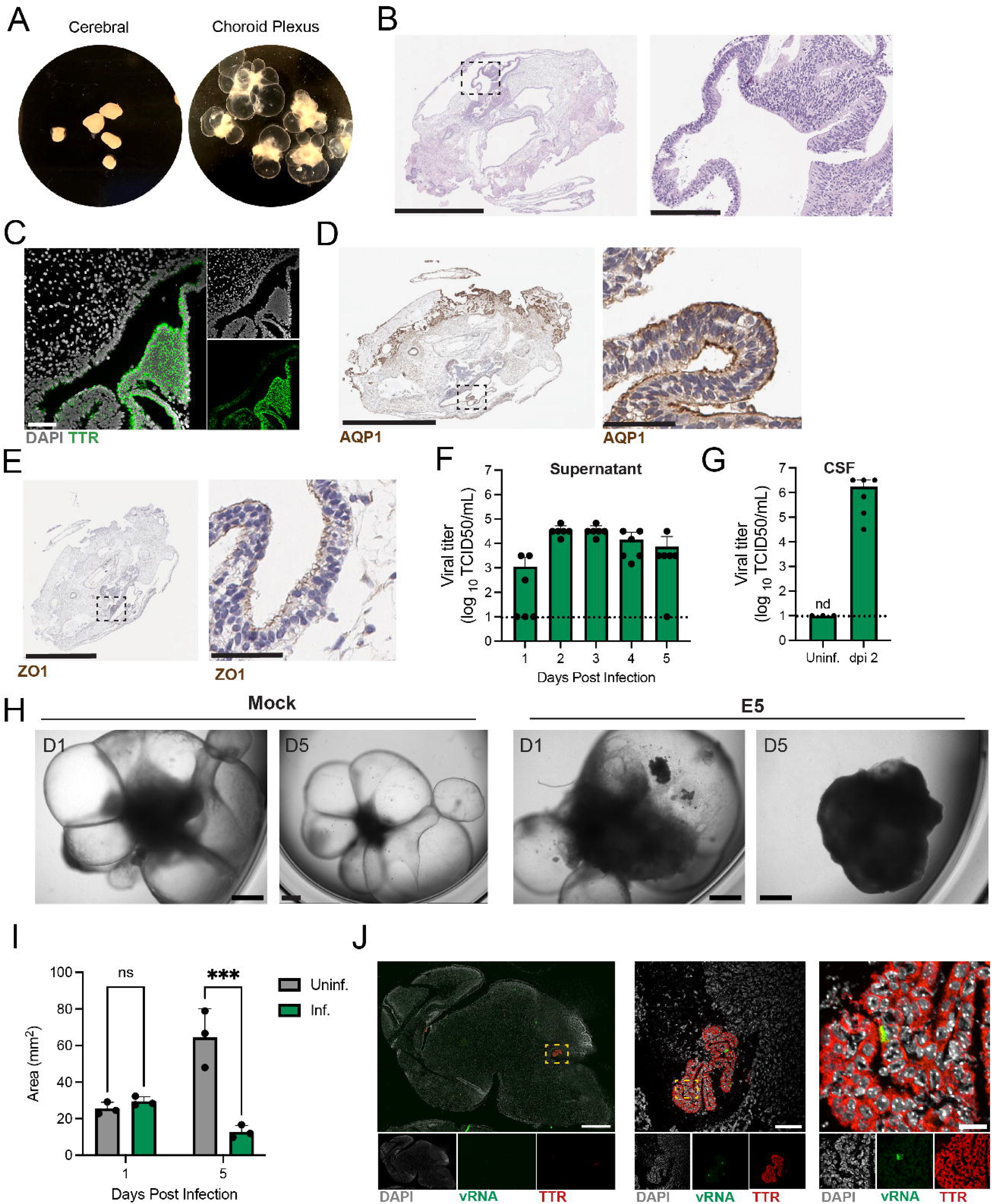
Echovirus infection of choroid plexus epithelium *in vitro* and *in vivo*. All choroid plexus organoids used were 55±5 days old **(A-I)**. **(A)** Brightfield images of 70-day-old cerebral organoids (left) and ChP organoids (right) in culture. **(B)** H&E staining of a ChP organoid showing epithelial cells lining a cystic structure; the dashed box indicates the region shown at higher magnification on the right. **(C)** RNAscope staining of a ChP organoid for transthyretin (*TTR*; green) and DAPI (gray). **(D)** Immunohistochemical staining of a ChP organoid for aquaporin-1 (AQP1); the dashed box indicates the region shown at higher magnification. **(E)** Immunohistochemical staining of a ChP organoid for ZO-1; the dashed box indicates the region shown at higher magnification. **(F)** Viral titers in supernatants from ChP organoids infected with 200 PFU of E5 virus, measured by TCID50 assay and shown as log_10_ TCID50 per milliliter; the dotted line indicates the limit of detection, with data shown as mean ± standard deviation and individual organoids represented as points. **(G)** Viral titers in CSF-like fluid collected from ChP organoids, measured by TCID50 assay and shown as log_10_ TCID50 per milliliter; the dotted line indicates the limit of detection, with data shown as mean ± standard deviation and individual organoids represented as points. **(H)** Brightfield images of mock- and E5-infected ChP organoids at one and five days post infection. **(I)** Quantification of organoid area at one and five days post infection; data are shown as mean ± standard deviation with individual organoids represented as points. Statistical significance was determined by two-way ANOVA with repeated measures (p < 0.0001). **(J)** RNAscope staining for viral RNA (vRNA; green) and *Ttr* (red) in a representative brain section from a Tg32 mouse at two days post infection; the dashed box indicates the region shown at higher magnification on the right. Scale bars: 2 mm and 200 µm (A, B), 100 µm (C), 2 mm and 50 µm (D, E), 1 mm (H), and 1 mm, 100 µm, and 20 µm (J).

To assess echovirus infection and replication dynamics, ChP organoids were infected with 200 PFU of E5 and viral replication was monitored over a five-day time course. Infection experiments were performed using two independent organoid batches, with three organoids per condition in each batch. High titers of infectious virus were detected in the supernatant throughout the time course, indicating productive infection (**Figure 2F**). Because echovirus RNA is frequently detected in the CSF of patients with aseptic meningitis ^14^, we next asked whether infectious virus could access the CSF compartment in this organoid model. CSF-like fluid was collected from ChP organoids from three independent batches at two days post infection, and high titers of infectious virus were detected, indicating that E5 is present within the CSF compartment in this system (**Figure 2G**). In parallel, pronounced cytopathic effects were observed in infected ChP organoids over the course of infection. By five days post infection, the large, fluid-filled cystic structures had collapsed, and infected organoids were markedly smaller than uninfected controls, consistent with epithelial damage following E5 infection (**Figure 2H-I**). Together, these findings establish that ChP organoids provide a robust *in vitro* platform to model viral infection and epithelial damage at the blood-CSF interface.

To extend these observations *in vivo*, we employed a mouse model of echovirus infection using transgenic animals that express human *FCGRT* under the control of its endogenous promoter (hFcRnTg32 mice, hereafter referred to as Tg32)^31,32^. Using this system, we developed an intracranial infection model in which virus is introduced directly into the right cerebral hemisphere, enabling seeding of infection within the CNS. Given the age-dependent severity of echovirus disease, neonatal Tg32 mice at postnatal day three were inoculated with 200 PFU of E5, and brains were harvested at two days post infection. RNAscope analysis for viral RNA together with markers for the choroid plexus epithelium (*Ttr*), neurons (*Map2*), astrocytes (*Gfap*), and neural progenitor cells (*Sox2*) revealed that infection was highly restricted to the ChP epithelium (**Figure 2J, Supplemental Figure 2A-B**). Comparable levels of infection were observed across ChP regions in different ventricles (**Supplemental Figure 2C-D**), including the distant fourth ventricle, indicating that susceptibility is not regionally restricted and that viral spread is not limited by the location of inoculation.

### The Choroid Plexus Epithelium Engages a Type III Interferon Antiviral Program

To define the immunological response of the ChP epithelium to viral infection, we performed multianalyte Luminex profiling of 74 cytokines and chemokines in supernatants collected from infected ChP organoids at 1-, 3-, and 5-days post infection. Experiments were conducted using two independent organoid batches, with three organoids per condition in each batch. ChP organoids mounted a robust inflammatory response to infection, with more than 40 cytokines and chemokines exhibiting at least a two-fold increase at five days post infection compared with uninfected controls (**Figure 3A**, **Supplemental Figure 3A-C**). Among the most strongly induced analytes were type III IFNs, and to a lesser extent type I IFNs, with IFNλ-2/3 showing the highest induction of any analyte measured (**Figure 3B-C**). Because we detected infectious E5 in ChP organoid-derived CSF, we next asked whether IFNs produced by the choroid plexus epithelium are released into the CSF compartment. Luminex analysis of CSF-like fluid collected from infected organoids at 2- and 3-days post infection revealed high levels of both type I and type III IFNs, indicating that the ChP epithelium actively secretes these cytokines into the CSF during infection (**Figure 3C**). Finally, we asked if the type III IFN response we observed is specific to echovirus infection or if it is a more generalized antiviral response of the ChP epithelium. Quantitative PCR measurements for IFN expression in ChP organoids stimulated with the synthetic viral dsRNA mimic, poly(I:C), showed strong induction of both type I and type III IFNs compared to unstimulated controls (**Figure 3D**). Together, these data establish that type III IFNs are a prominent component of the ChP epithelial response to echovirus infections, and that they may play a broader role in the antiviral defenses of the ChP, however, responses to other viruses may vary according to tropism and immune-evasion mechanisms.

**Figure 3.**
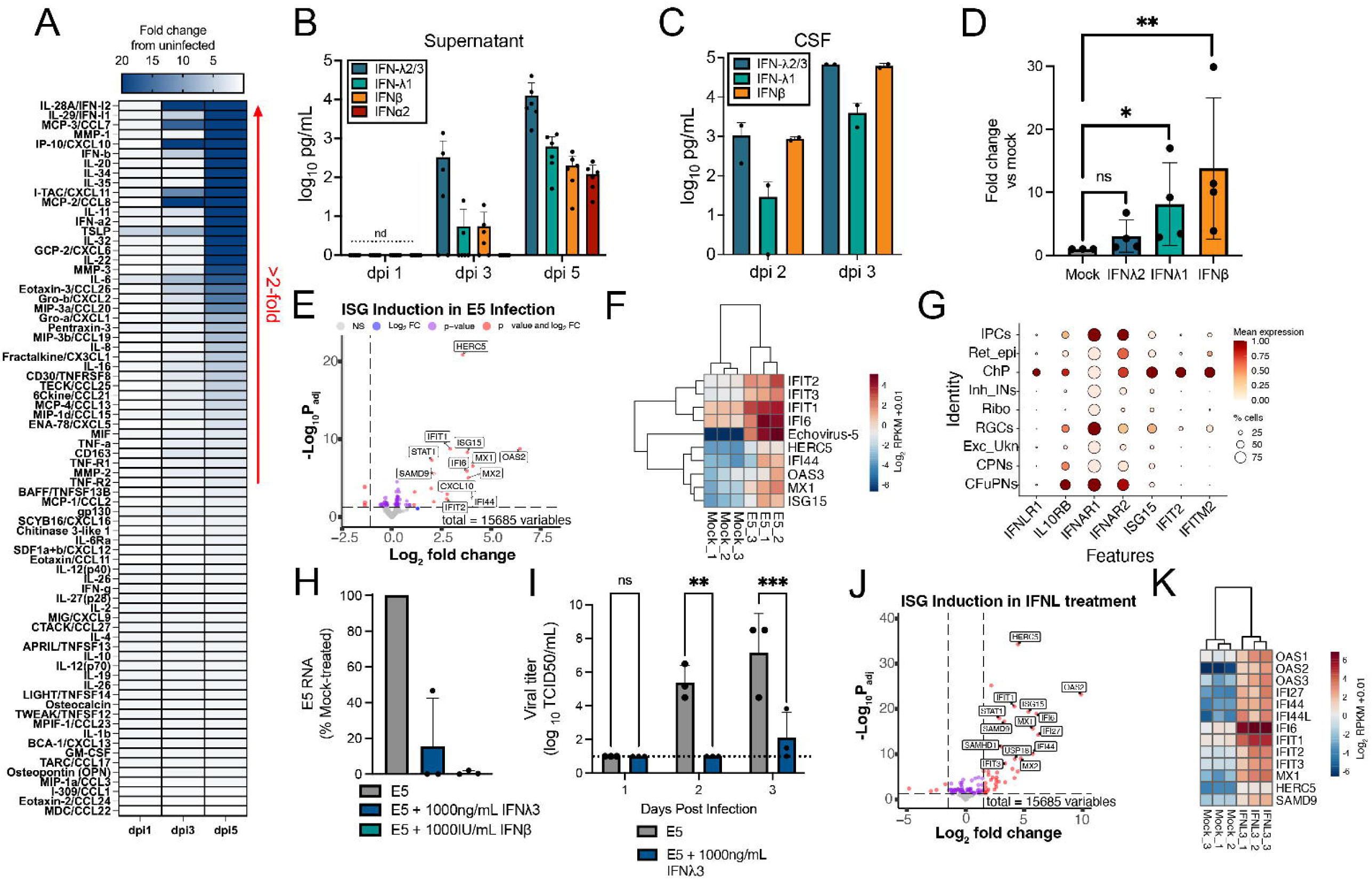
The choroid plexus epithelium mounts a type III IFN driven antiviral program. All choroid plexus organoids used were 55±5 days old. **(A)** Heatmap showing changes in cytokine and chemokine levels in supernatants collected from ChP organoids infected with 200 PFU of E5 at 1, 3, and 5 days post infection, displayed as the average fold change relative to uninfected controls (n = 6). **(B)** Bar plot showing concentrations (log_10_ pg/mL) of type III interferons (IFN-λ2/3 and IFN-λ1) and type I interferons (IFN-β and IFN-α2) in supernatants from infected ChP organoids at 1, 3, and 5 days post infection; data are shown as mean ± standard deviation with individual organoids represented as points. **(C)** Bar plot showing concentrations (log_10_ pg/mL) of type III interferons (IFN-λ2/3 and IFN-λ1) and type I interferons (IFN-β) in CSF-like fluid collected from ChP organoids at 2 and 3 days post infection; data are shown as mean ± standard deviation with individual organoids represented as points. **(D)** ChP organoids were stimulated with 10 μg/mL of poly(I:C) overnight. Bar plot showing the induction of IFNλ2, IFNλ1, and IFNβ as a fold change compared to untreated controls; data are shown as mean ± standard deviation with individual organoids represented as points. Statistical significance determined by unpaired t test using ΔCt values (* p = 0.0171, ** p = 0.0065). **(E)** Volcano plot showing differential gene expression in ChP organoids at three days post infection compared with mock-infected controls (n = 3). **(F)** Heatmap showing the top upregulated interferon-stimulated genes (ISGs) in ChP organoids at three days post infection (n = 3). **(G)** Dot plot from scRNAseq of infected cerebral organoids showing expression of type III IFN receptor subunits (*IFNLR1* and *IL10RB*), type I IFN receptor subunits (*IFNAR1* and *IFNAR2*), and selected ISGs (*ISG15*, *IFIT2*, and *IFITM2*) across annotated cell clusters; values are scaled such that 1 (red) represents the highest normalized expression and 0 (white) represents the lowest, and dot size indicates the percentage of cells within each cluster expressing the indicated gene. **(H-K)** ChP organoids were pretreated with 0, 1,000 ng/mL recombinant human IFN-λ3, or 1,000 IU/mL of recombinant human IFNβ prior to infection with 200 PFU of E5. **(H)** Bar plot showing E5 RNA levels measured by quantitative PCR in ChP organoids following IFN pretreatment; data are shown as mean ± standard deviation with individual organoids represented as points. **(I)** Viral titers in supernatants from ChP organoids pretreated with 1,000 ng/mL of recombinant human IFNλ3 prior to infection with 200 PFU of E5 virus, measured by TCID50 assay and shown as log_10_ TCID50 per milliliter; the dotted line indicates the limit of detection, with data shown as mean ± standard deviation and individual organoids represented as points. Statistical significance was determined by two-way ANOVA with repeated measures (** p = 0.0023, *** p = 0.0007). **(J)** Volcano plot showing differential gene expression in uninfected ChP organoids treated with 1,000 ng/mL IFN-λ3 (n = 3). **(K)** Heatmap showing the top upregulated ISGs in uninfected ChP organoids treated with 1,000 ng/mL IFN-λ3 (n = 3). For volcano plots, pink circles indicate significantly differentially expressed genes (log₂ fold change > 1; adjusted p value < 0.05). For heatmaps, red indicates higher expression and blue indicates lower expression.

To define the transcriptional programs altered by echovirus infection, we performed bulk RNA sequencing on infected and uninfected ChP organoids. Reads were aligned to both the human and viral genomes, and differential gene expression analysis was carried out using DESeq2 (**Supplemental Table 2**). Differentially expressed genes were defined as those with a log2 fold change greater than 1 and an adjusted p value less than 0.05 (**Supplemental Table 3**). Among genes upregulated in infected organoids, a large proportion corresponded to canonical interferon-stimulated genes (ISGs), including *IFIT1*, *ISG15*, and *HERC5*, indicating that infection induces a robust IFN driven antiviral transcriptional program in the ChP epithelium (**Figure 3E-F**).

We next examined whether the ChP epithelium is positioned to respond to type III interferons. Although type III IFN signaling is largely restricted to epithelial cells and select immune populations, its presence at the ChP has not been well characterized. To address this, we returned to our scRNA-seq dataset from cerebral organoids and examined expression of IFN receptors across cell populations. While several clusters expressed the type I IFN receptor subunits, *IFNAR1* and *IFNAR2*, only the ChP epithelial cell cluster expressed both components of the type III IFN receptor, *IFNLR1* and *IL10RB* (**Figure 3G**). Consistent with this, analysis of ISG expression revealed that ChP epithelial cells uniquely exhibited high levels of multiple ISGs during infection (**Figure 3G**).

To test whether type III IFNs are sufficient to confer antiviral protection, we pretreated ChP organoids with either 1,000 ng per mL of recombinant human IFNλ3 or 1,000 IU per mL of recombinant human IFNβ prior to infection and assessed viral burden. Pretreatment with IFNβ resulted in a marked reduction in echovirus RNA levels compared with untreated controls, as measured by quantitative PCR, matching previous reports of the antiviral capabilities of type I IFNs at the choroid plexus ^21,33^. Pretreatment of ChP organoids with IFNλ3 also resulted in an approximately 85 percent reduction in echovirus RNA (**Figure 3H**). In addition to reduced viral RNA, pretreatment with IFNλ3 also resulted in a significant reduction in viral titers at days two and three post infection as measured by TCID50 (**Figure 3I)**. The antiviral capacity of both IFNβ and IFNλ3 also extended to post infection treatment timepoints where they resulted in a reduction in E5 RNA compared to untreated controls (**Supplemental Figure 3D)**. Finally, to define the transcriptional response to type III IFN in the absence of infection, we performed bulk RNA sequencing on uninfected organoids treated with IFNλ3 (**Supplemental Table 2**). More than 70 percent of upregulated genes were ISGs, and the resulting signature closely mirrored the antiviral program induced by echovirus infection and IFNβ treatment (**Figure 3J-K**, **Supplemental Figure 3E, Supplemental Table 4**). Together, these findings demonstrate that the ChP epithelium not only produces type III IFNs in response to viral infection but is also uniquely poised to respond to this pathway, mounting a potent IFN driven antiviral program.

### Type I Interferon Signaling Restricts Echovirus Infection in the Choroid Plexus

To define the roles of type I and type III IFNs in antiviral defense at the ChP *in vivo*, we used previously established genetically tractable mouse models that enable selective dissection of IFN signaling during echovirus infection ^24,34,35^. To do this, we used Tg32 mice, which express the human neonatal Fc receptor required for echovirus entry, together with matched lines lacking the type I interferon receptor (Tg32*^Ifnar1-/-^*), the type III interferon receptor (Tg32*^Ifnlr1-/-^*), or both receptors (Tg32*^Ifnar1-/-Ifnlr1-/-^*). Neonatal mice from all four genotypes were inoculated intracranially with 200 PFU of E5 and monitored for survival. Strikingly, all mice deficient in type I IFN signaling (Tg32*^Ifnar1^*^⁻/⁻^ and Tg32*^Ifnar1^*^⁻/⁻^*^Ifnlr1^*^⁻/⁻)^ succumbed to infection by three days post infection, whereas mice with intact type I IFN responses (Tg32 and Tg32*^Ifnlr1^*^⁻/⁻^) survived beyond this acute phase (**Figure 4A**). These survival differences were mirrored by viral burden in the brains at two days post infection. Although all genotypes harbored detectable virus, Tg32*^Ifnar1^*^⁻/⁻^ and Tg32*^Ifnar1^*^⁻/⁻^*^Ifnlr1^*^⁻/⁻^ mice exhibited significantly higher viral titers than Tg32 and Tg32*^Ifnlr1^*^⁻/⁻^ mice (**Figure 4B**).

**Figure 4.**
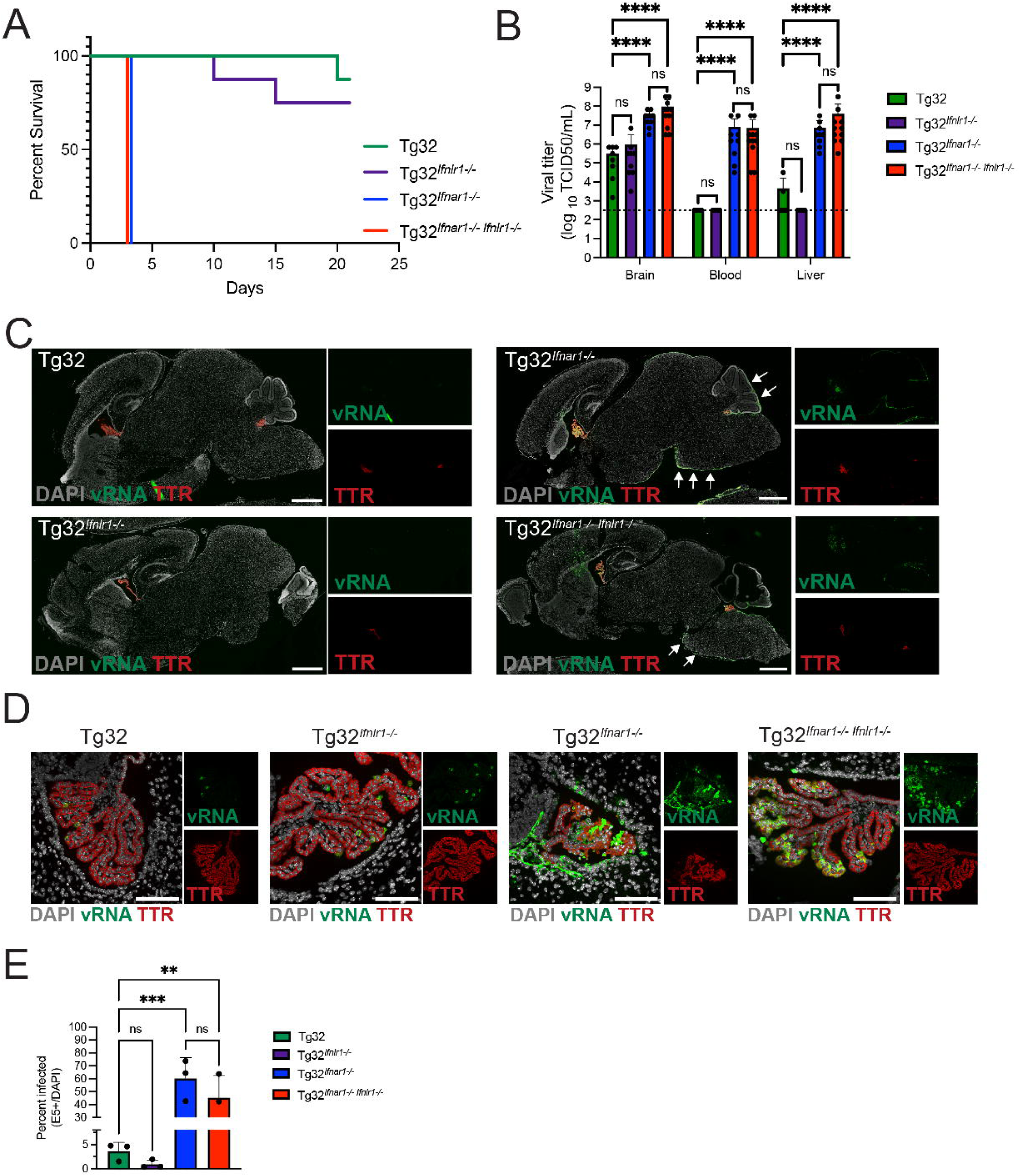
Type I interferon signaling limits echovirus infection in the choroid plexus. **(A-E)** All four genotypes (Tg32, Tg32*^Ifnlr1-/-^*, Tg32*^Ifnar1-/-^*, and Tg32*^Ifnar1-/-Ifnlr1-/-^*) were intracranially inoculated with 200 PFU of E5 at postnatal day three. **(A)** Survival curves for each genotype; n = 8 pups per group. **(B)** Viral titers in brain, blood, and liver at two days post infection, measured by TCID50 assay and shown as log_10_ TCID50 per milliliter; the dotted line indicates the limit of detection. Data are shown as mean ± standard deviation with individual animals represented as points. Statistical significance was assessed by one-way ANOVA (p < 0.0001). **(C)** Tile-scanned whole-brain sections at two days post infection stained by RNAscope for viral RNA (vRNA; green) and transthyretin (*Ttr*; red); white arrows indicate vRNA signal in the meninges. **(D)** RNAscope staining for vRNA (green) and *Ttr* (red) in representative choroid plexus regions from each genotype. **(E)** Quantification of the percentage of infected choroid plexus epithelial cells, shown as mean ± standard deviation with individual mice represented as points. Statistical significance was determined by an ordinary one-way ANOVA with multiple comparisons (** p = 0.0047, *** p = 0.0005). Scale bars: 1 mm (C) and 100 µm (D).

To assess whether loss of type I IFN signaling also promoted systemic dissemination, we quantified viral titers in the blood and liver, a known site of echovirus replication. Mice lacking the type I IFN receptor showed markedly elevated viral loads in both compartments compared with Tg32 and Tg32*^Ifnlr1^*^⁻/⁻^ mice (**Figure 4B**). Notably, none of the Tg32*^Ifnlr1^*^⁻/⁻^ mice had detectable virus in the blood or liver, and only a minority of Tg32 mice showed low level liver infection. Viral titers did not differ by sex (**Supplemental Figure 4A**). Together, these data demonstrate that type I IFN signaling, but not type III IFN signaling, is indispensable for preventing lethal disease and restricting systemic spread of echovirus *in vivo*.

Because bulk viral titers from whole-brain homogenates cannot resolve infection dynamics at the ChP, we performed RNAscope analysis for viral RNA together with *Ttr* on brain sections collected at two days post infection. This approach enabled spatially resolved assessment of viral replication across the brain and ventricular system. Whole brain imaging revealed that in Tg32 and Tg32*^Ifnlr1^*^⁻/⁻^ mice, infection was sparse and exclusively confined to the ChP epithelium (**Figure 4C-D**). In contrast, mice lacking the type I IFN receptor exhibited extensive viral signal within the ChP as well as in the meninges (**Figure 4C-D**), consistent with prior work demonstrating a central role for type I IFNs in controlling echovirus infection at meningeal surfaces following intraperitoneal inoculation ^36^. Image analysis evaluating the percentage of infected cells within the ChP across all four genotypes revealed that mice lacking the type I IFN receptor, Tg32*^Ifnar1^*^⁻/⁻^ and Tg32*^Ifnar1^*^⁻/⁻^*^Ifnlr1^*^⁻/⁻^, had significantly higher viral burden in the choroid plexus compared to Tg32 mice. However, there was not a significant difference between either the Tg32 and Tg32*^Ifnlr1-/-^* mice or the Tg32*^Ifnar1^*^⁻/⁻^ and Tg32*^Ifnar1^*^⁻/⁻^*^Ifnlr1^*^⁻/⁻^ mice (**Figure 4E**). Together, these data indicate that type I IFN signaling is required to limit echovirus replication at the ChP and to restrict viral spread within the CNS.

### Type I and Type III Interferons Drive ISG Expression in the Choroid Plexus Epithelium *In Vivo*

To determine whether the ChP epithelium can respond to type III IFNs *in vivo*, we reanalyzed a published single nucleus transcriptomic atlas of the mouse ChP comprising 83,040 cells from the lateral, third, and fourth ventricles of embryonic, adult, and aged animals ^37^. Because embryonic cells were transcriptionally distinct from postnatal populations, our analysis focused on adult and aged mice. Cells passing quality control were visualized using UMAP projections (**Figure 5A**), and cell populations were annotated based on established marker gene expression. This analysis identified six epithelial clusters, designated Epi-1 through Epi-6, each expressing canonical ChP epithelial markers including *Ttr*, *Enpp2*, *Aqp1*, *Atp1b1*, *Folr1*, and *Slc4a10* (**Figure 5B**). In addition, we identified multiple non-epithelial populations, including mesenchymal fibroblasts marked by *Col1a1*, *Col1a2*, and *Lum*, endothelial cells expressing *Cldn5*, *Pecam1*, and *Esam*, neurons expressing *Dcx*, *Rbfox3*, and *Neurod6*, and macrophages expressing *Ptprc*, *Cd74*, and *Cd68*. These populations were consistent with those described in the original dataset. We next examined expression of receptor subunits and signaling components required for responsiveness to type I and type III IFNs. All identified cell types expressed both subunits of the type I interferon receptor, *Ifnar1* and *Ifnar2*. In contrast, expression of the type III IFN receptor was restricted to epithelial clusters and macrophages, as only these populations expressed both *Ifnlr1* and *Il10rb* (**Figure 5C**). All cell types expressed the downstream signaling machinery required for IFN responses (**Figure 5C**). Merged feature plots demonstrated strong colocalization of *Ifnlr1* with the ChP epithelial marker *Aqp1* in epithelial clusters, whereas *Ifnlr1* expression in macrophages occurred in the absence of epithelial markers (**Figure 5D**). Together, these data establish that, in the mouse ChP, epithelial cells are uniquely positioned to respond to type III IFNs.

**Figure 5.**
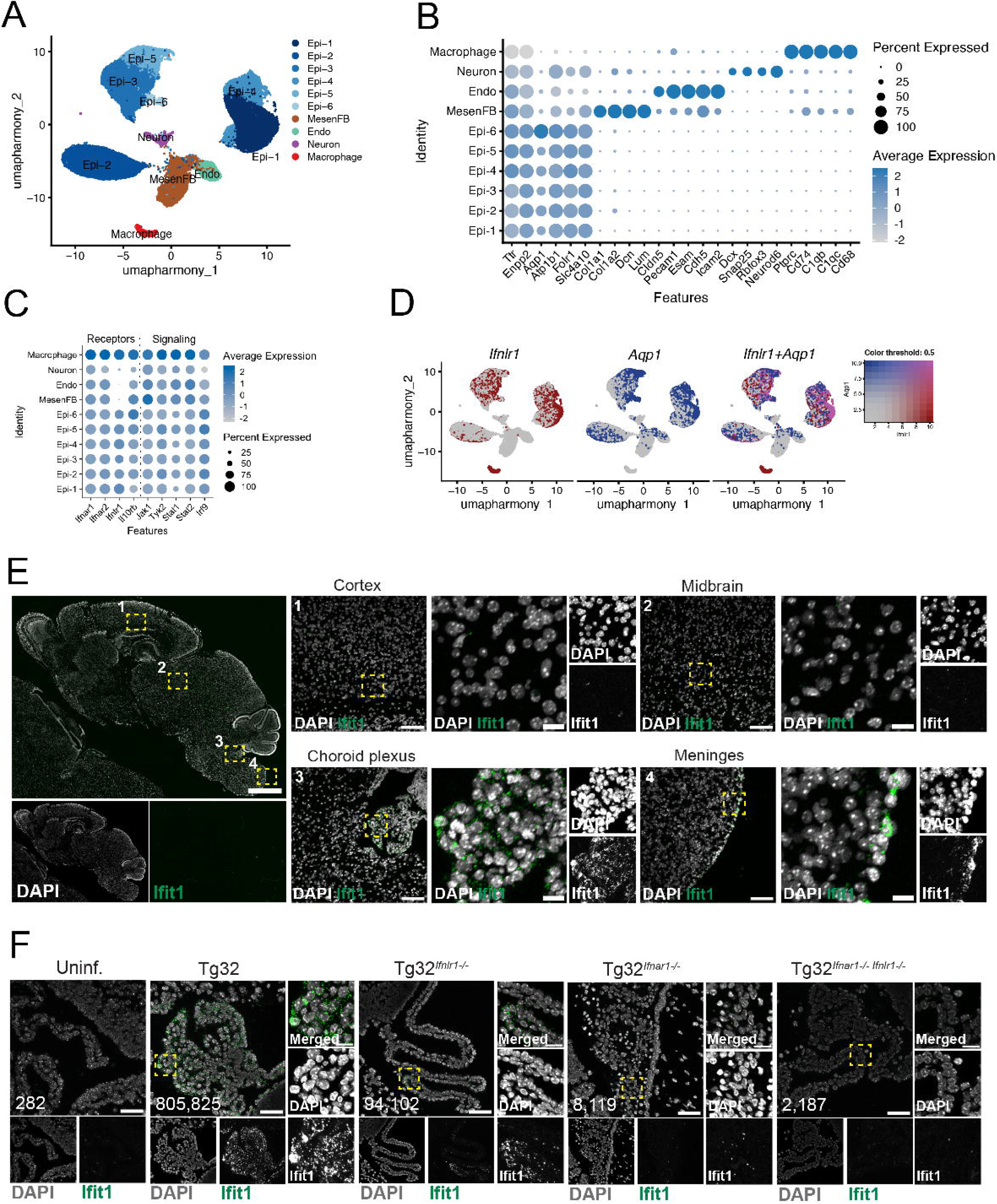
Interferon Receptor Expression and ISG Induction in the Choroid Plexus Epithelium *In Vivo*. **(A-D)** Reanalysis of a published single-nucleus RNA-sequencing dataset from adult and aged mouse choroid plexus samples. **(A)** UMAP showing clustering of choroid plexus epithelial cells (Epi-1–Epi-6), mesenchymal fibroblasts (MesenFB), endothelial cells (Endo), neurons, and macrophages. **(B)** Dot plot showing expression of selected marker genes for each annotated cell type. **(C)** Dot plot showing expression of type I interferon receptor subunits (*Ifnar1*, *Ifnar2*), type III interferon receptor subunits (*Ifnlr1*, *Il10rb*), and downstream signaling components across clusters. **(D)** Merged feature plot showing expression of *Ifnlr1* (red) and the choroid plexus epithelial marker *Aqp1* (blue); purple indicates cells expressing both features. **(E–F)** Tg32, Tg32Ifnlr1⁻/⁻, Tg32Ifnar1⁻/⁻, and Tg32Ifnar1⁻/⁻Ifnlr1⁻/⁻ mice were intracranially inoculated with 200 PFU of E5 at postnatal day three. **(E)** Tile-scanned whole-brain section from a Tg32 mouse at two days post infection stained by RNAscope for DAPI (gray) and *Ifit1* (green). Numbered dashed boxes indicate regions shown at higher magnification, including cortex (1), midbrain (2), choroid plexus (3), and meninges (4). **(F)** RNAscope staining for DAPI (gray) and *Ifit1* (green) in representative choroid plexus regions from each genotype and uninfected controls; dashed boxes indicate regions shown at higher magnification. Values in the lower left corner indicate total *Ifit1* signal intensity. High-magnification images in (E–F) are pseudocolored white and brightness adjusted to enhance visibility. Scale bars: 1 mm, 100 µm, and 20 µm (E); 50 µm and 20 µm (F). For dot plots, values are scaled such that 1 (blue) represents the highest normalized expression and 0 (white) represents the lowest normalized expression; dot size indicates the percentage of cells within each cluster expressing the indicated gene.

Having defined the receptor landscape, we next determined whether the ChP epithelium mounts an ISG response during infection *in vivo*. To address this, we performed RNAscope staining for the canonical ISG *Ifit1* on tile-scanned whole-brain sections from Tg32, Tg32*^Ifnlr1^*^⁻/⁻^, Tg32*^Ifnar1^*^⁻/⁻^, and Tg32*^Ifnar1^*^⁻/⁻^*^Ifnlr1^*^⁻/⁻^ mice at two days post infection, alongside uninfected controls. Whole-brain imaging revealed that ISG induction was largely confined to the ChP epithelium and was not detected in the surrounding brain parenchyma, highlighting the spatial restriction of IFN responses at this barrier interface (**Figure 5E**). Robust *Ifit1* expression was detected in the ChP epithelium of Tg32 mice, with reduced signal in Tg32*^Ifnlr1^*^⁻/⁻^ animals (**Figure 5F**). In contrast, *Ifit1* expression was virtually undetectable in the ChP epithelium of Tg32*^Ifnar1^*^⁻/⁻^ and Tg32*^Ifnar1^*^⁻/⁻^*^Ifnlr1^*^⁻/⁻^ mice and was comparable to levels observed in uninfected controls (**Figure 5F**). Together, these data show that *in vivo* the ChP epithelium expresses the receptor components required for type III IFN signaling and exhibits a localized ISG response during viral infection at the blood-CSF interface.

### Type III Interferons Promote Epithelial Barrier Injury at the Choroid Plexus During Infection

To assess the impact of echovirus infection on tissue integrity at the ChP *in vivo*, we first performed histopathological analysis across all four mouse genotypes. Hematoxylin and eosin (H&E) staining revealed minimal structural disruption of the ChP epithelium in Tg32 and Tg32*^Ifnlr1^*^⁻/⁻^ mice, consistent with their lower levels of infection (**Figure 6A**). In contrast, Tg32*^Ifnar1⁻/⁻^* mice exhibited pronounced epithelial pathology, characterized by epithelial disorganization and tissue damage, whereas Tg32*^Ifnar1⁻/⁻Ifnlr1⁻/⁻^* mice displayed minimal ChP pathology despite comparable viral burdens (**Figure 6A**). This unexpected dissociation between viral load and tissue injury suggested that IFN signaling pathways differentially influence epithelial integrity at the ChP during infection.

**Figure 6.**
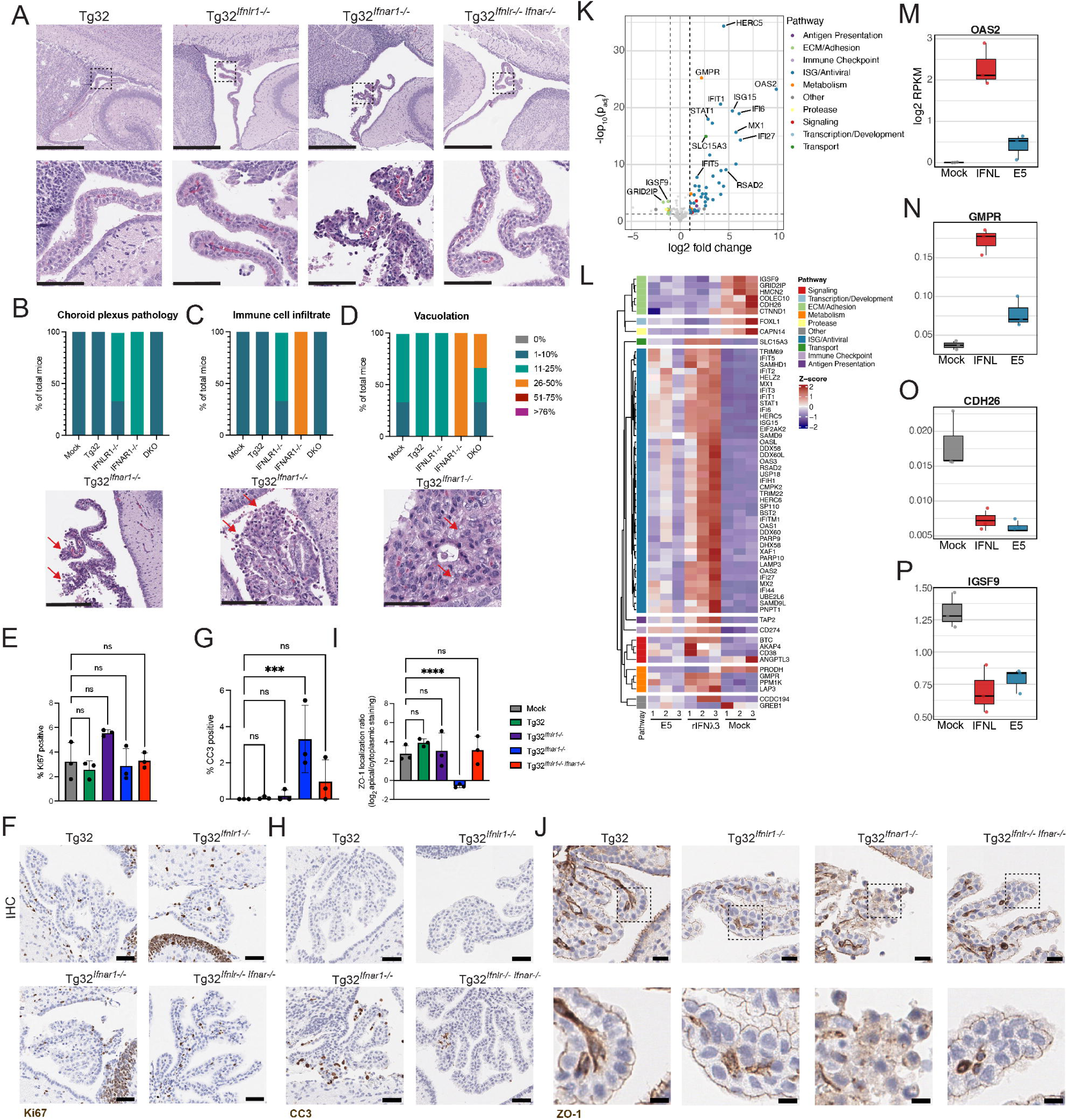
Type III interferon-associated epithelial barrier injury at the choroid plexus during infection. **(A-J)** Tg32, Tg32*^Ifnlr1-/-^*, Tg32*^Ifnar1-/-^*, and Tg32*^Ifnar1-/-Ifnlr1-/-^* mice were intracranially inoculated with 200 PFU of E5 at postnatal day three, and brains were harvested at two days post infection. **(A)** Representative hematoxylin and eosin (H&E)-stained sections showing the choroid plexus across all four genotypes; the dotted box indicates the region shown at higher magnification below. **(B–D)** Blinded pathological scoring of H&E-stained sections for **(B)** choroid plexus epithelial pathology, **(C)** immune cell infiltration, and **(D)** epithelial vacuolation across all four genotypes, with representative images shown below (n = 3). The scale at right indicates the percentage of tissue affected; red arrows denote representative pathological features. **(E-H)** Quantification of immunohistochemical (IHC) staining for (E-F) the cell proliferation marker Ki67 and (G-H) the apoptosis marker CC3 within the choroid plexus with representative images. Data are shown as mean ± standard deviation with individual mice represented as points. Statistical significance was determined by two-way ANOVA with multiple comparisons (*** p = 0.0002). **(I-J)** Quantification of IHC staining for the tight junction protein ZO-1 in the choroid plexus across all four genotypes. Quantification is represented as the ratio of apical staining intensity to cytoplasmic staining intensity with representative images. Data are shown as mean ± standard deviation with individual mice represented as points. Statistical significance was determined using two-way ANOVA with multiple comparisons (**** p < 0.0001). **(K-P)** Choroid plexus organoids were pretreated with 0 or 1,000 ng/mL recombinant human IFN-λ3, or infected with 200 PFU of E5; RNA was collected at three days post infection/treatment (n = 3). **(K)** Volcano plot showing differentially expressed genes in IFN-λ3–treated choroid plexus organoids, with point colors indicating associated Gene Ontology (GO) pathways (key at right). **(L)** Heatmap comparing differentially expressed genes across E5-infected, IFN-λ3–treated, and untreated choroid plexus organoids; colors at left denote associated GO pathways (key at right), with red indicating higher expression and blue indicating lower expression. **(M-P)** Expression changes of selected individual genes across the three conditions. Scale bars: 500 µm and 100 µm (A), 200 µm (B), 100 µm (C), 50 µm (D-H), and 50 µm and 10 µm (J).

To quantify these observations, a board-certified pathologist performed blinded evaluation of the ChP from brain sections from three mice per genotype and uninfected controls, scoring ChP pathology, immune cell infiltration, and epithelial vacuolation. Across all parameters, Tg32*^Ifnar1^*^⁻/⁻^ mice displayed the most severe pathology (**Figure 6B-D, Supplemental Figure 5A**), including frequent epithelial discontinuities, marked inflammatory cell infiltration, and extensive cytoplasmic vacuolation, a feature associated with virus-induced epithelial stress. In contrast, Tg32*^Ifnar1^*^⁻/⁻^*^Ifnlr1^*^⁻/⁻^ mice exhibited significantly reduced tissue damage despite similar levels of viral replication. These findings indicate that epithelial injury at the ChP is not solely determined by viral burden.

Type III IFNs have been reported to impair epithelial repair and barrier integrity in peripheral tissues, including the lung and intestine, during infection and chronic inflammation through their antiproliferative and pro-apoptotic capabilities ^25,38,39^. We therefore asked whether a similar mechanism contributes to ChP pathology in this setting. To determine if the tissue damage seen in the Tg32*^Ifnar1-/-^* mice was driven by either an increase in cell death or a decrease in cell proliferation in ChP epithelial cells, we performed immunohistochemical staining for an apoptotic marker, cleaved caspase 3 (CC3), and a cell proliferation marker, Ki67. While there was no significant difference in Ki67 staining in the ChP epithelium across our mice (**Figure 6E-F, Supplemental Figure 5B**), we did observe an increase in CC3 staining in the epithelium of Tg32*^Ifnar1-/-^* mice (**Figure 6G-H, Supplemental Figure 5C**). A similar increase in cell death was reported in both lung and intestinal epithelial cells following viral recognition and subsequent type III IFN production, indicating a similar mechanism may play a role in the ChP ^25,38,39^.

To determine if this tissue damage also impacted the junctional organization of the ChP epithelium, we performed immunohistochemical staining for the junctional protein ZO-1. In Tg32 and Tg32*^Ifnlr1^*^⁻/⁻^ mice, which harbored lower viral loads, ZO-1 localization remained sharply delineated along epithelial borders, comparable to uninfected controls. In contrast, among the genotypes with high viral burdens, only Tg32*^Ifnar1^*^⁻/⁻^ mice exhibited marked disruption of ZO-1 localization with staining becoming diffuse and cytoplasmic, whereas Tg32*^Ifnar1^*^⁻/⁻^*^Ifnlr1^*^⁻/⁻^ mice retained intact junctional architecture (**Figure 6I-J, Supplemental Figure 5D**). We also performed immunohistochemical staining for the broad macrophage marker F4/80 to determine if macrophages contributed to the increase in immune cell infiltration seen in the ChP of Tg32*^Ifnar1^*^⁻/⁻^ mice. There was no significant difference in the number of macrophages at the ChP epithelium across genotypes, indicating that macrophage infiltration levels are not influencing the increased pathology seen at this acute timepoint (**Supplemental Figure 5E-F)**. These data indicate that epithelial injury and tight junction disorganization reflect a specific effect of type III IFN signaling rather than infection severity alone.

To define molecular programs associated with this phenotype, we returned to bulk transcriptomic analysis of human ChP organoids treated with recombinant IFNλ3. Differential expression and pathway analyses revealed coordinated suppression of gene networks involved in epithelial barrier maintenance, including extracellular matrix organization, cell–cell adhesion, and epithelial differentiation (**Figure 6K-L**, **Supplemental Table 4**). Among the most significantly downregulated genes were those encoding key structural and adhesion components of epithelial barriers, including *CDH26*, *IGSF9*, and *CAPN14*, as well as regulators of epithelial identity and maturation such as *FOXL1*. In parallel, IFNλ3 treatment reduced expression of genes implicated in extracellular matrix organization, including *HMCN2* and *COLEC10*. Conversely, canonical antiviral effectors such as *OAS2* and *IFIT1* were among the most strongly induced transcripts, underscoring a transcriptional trade-off in which sustained antiviral programming occurs at the expense of barrier repair and epithelial homeostasis (**Figure 6M-P**). Notably, this transcriptional signature mirrors gene programs previously linked to defective epithelial regeneration in other barrier tissues exposed to prolonged type III IFN signaling ^25,39^, suggesting that IFNλ imposes a conserved barrier-disruptive state across epithelial interfaces.

### Type III Interferon Signaling Promotes Ventricular Enlargement and Ependymal Cell Loss Following Viral Clearance

Aseptic meningitis in early life is frequently associated with long-term neurological sequelae, yet the mechanisms linking acute infection to chronic structural and developmental abnormalities remain poorly defined ^5^. Having observed that type III IFN signaling compromised ChP epithelial integrity during acute echovirus infection, we next asked whether this pathway also contributes to pathological outcomes after viral clearance. To address this, we established a post-infection recovery model using Tg32 and Tg32*^Ifnlr1⁻/⁻^* mice, which survive the acute phase of infection. Neonatal mice were inoculated intracranially with 1,000 PFU of E5 and monitored through convalescence. Both genotypes fully cleared virus by fourteen days post infection (**Figure 7A**), allowing assessment of post-infectious sequelae independent of ongoing viral replication. Although both Tg32 and Tg32*^Ifnlr1⁻/⁻^* mice exhibited transient weight loss and a modest drop in survival during recovery (**Supplemental Figure 6A-B**), infected Tg32 mice displayed markedly more severe growth impairment than their *Ifnlr1*-deficient counterparts (**Figure 7B, Supplemental Figure 6C**). Given the link between echovirus infection and delayed developmental outcomes ^5^, we assessed the motor functions of long-term recovery mice using the surface righting reflex test. We found that infected Tg32 mice had a significantly impaired righting reflex time compared to uninfected controls, which was not observed in Tg32*^Ifnlr1-/-^* mice (**Figure 7C**). In addition, a subset of infected Tg32 mice developed overt cranial abnormalities, including head swelling and shortened snouts, phenotypes not observed in infected Tg32*^Ifnlr1⁻/⁻^* animals (**Supplemental Figure 6C**). While head-to-body ratios did not differ significantly between groups (**Figure 7D**), these gross morphological changes prompted us to examine ventricular architecture. Because hydrocephalus commonly arises from dysregulated CSF dynamics and manifests as ventriculomegaly ^40^, we quantified ventricular size using histological analysis. Strikingly, half of infected Tg32 mice exhibited marked ventricular enlargement, compared with only one infected Tg32*^Ifnlr1^*^⁻/⁻^ mouse (**Figure 7E-G**). Ventriculomegaly was defined as ventricular area exceeding twice the mean of uninfected controls. These findings demonstrate that type III IFN signaling promotes long-term structural abnormalities of the ventricular system following resolution of infection, and that these structural abnormalities may impact developmental milestones related to motor function and coordination.

**Figure 7.**
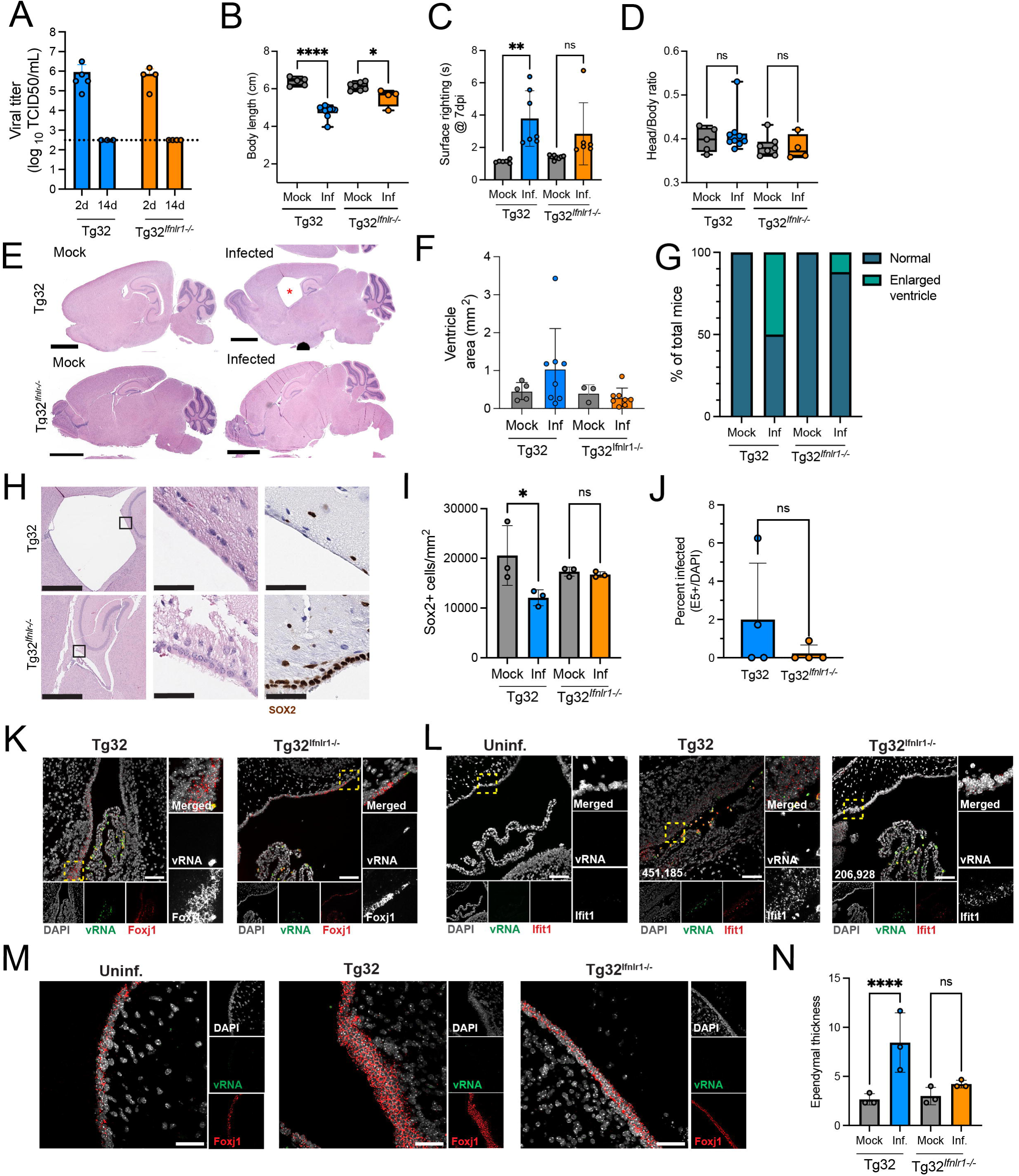
Type III interferon signaling is associated with ventricular enlargement and ependymal cell loss following viral clearance. **(A–N)** Tg32 and Tg32^Ifnlr1⁻/⁻^ mice were intracranially inoculated with 1,000 PFU of E5 at postnatal day three and monitored through viral clearance (∼17 days post infection). **(A)** Viral titers in Tg32 and Tg32^Ifnlr1⁻/⁻^ mice at 2 and 14 days post infection, measured by TCID50 assay and shown as log_10_ TCID50 per milliliter; the dotted line indicates the limit of detection. Data are shown as mean ± standard deviation with individual animals represented as points. **(B)** Body length of mock- and E5-infected Tg32 and Tg32^Ifnlr1⁻/⁻^ mice, shown as box-and-whisker plots with individual animals represented as points; statistical significance was assessed by unpaired t test (**** p < 0.0001; * p = 0.0169). **(C)** Surface righting reflex time at seven days post infection in mock- and E5-infected Tg32 and Tg32^Ifnlr1⁻/⁻^ mice. Data shown as mean ± standard deviation with individual animals represented as points; statistical significance was assessed by two-way ANOVA (** p = 0.006). **(D)** Head-to-body length ratio of mock- and E5-infected Tg32 and Tg32^Ifnlr1⁻/⁻^ mice; no significant differences were detected by unpaired t test. **(E)** Representative H&E-stained whole-brain sections from mock- and E5-infected Tg32 and Tg32^Ifnlr1⁻/⁻^ mice; the red asterisk indicates an enlarged ventricle. **(F)** Quantification of ventricular area in mock- and E5-infected Tg32 and Tg32^Ifnlr1⁻/⁻^ mice, shown as mean ± standard deviation with individual animals represented as points. **(G)** Percentage of mock- and E5-infected Tg32 and Tg32^Ifnlr1⁻/⁻^ mice exhibiting ventricular enlargement. **(H)** Representative H&E-stained sections and serial sections immunohistochemically stained for SOX2 from an E5-infected Tg32 mouse with ventricular enlargement and an E5-infected Tg32^Ifnlr1⁻/⁻^ mouse; the boxed region indicates the area shown at higher magnification. **(I)** Quantification of SOX2-positive cells lining the ventricular wall in mock- and E5-infected Tg32 and Tg32^Ifnlr1⁻/⁻^ mice. Data shown as mean ± standard deviation; statistical significance was assessed by unpaired t test (p = 0.0477). **(J)** Quantification of the percentage of infected ependymal cells, shown as mean ± standard deviation with individual ventricles represented as points; no significant differences were detected by unpaired t test. **(K)** Representative RNAscope images showing viral RNA (vRNA; green) and the ependymal marker *Foxj1* (red) in Tg32 and Tg32^Ifnlr1⁻/⁻^ mice at two days post infection; the dashed box indicates the region shown at higher magnification. **(L)** Representative RNAscope images showing vRNA (green) and *Ifit1* (red) in Tg32 and Tg32^Ifnlr1⁻/⁻^ mice at two days post infection; the dashed box indicates the region shown at higher magnification. Values in the lower left corner indicate total *Ifit1* signal intensity. **(M)** Representative RNAscope images showing vRNA (green) and *Foxj1* (red) in uninfected and E5-infected Tg32 and Tg32*^Ifnlr1-/-^* mice at two days post infection. **(N)** Quantification of the thickness of the ependymal cell layer in both uninfected and infected Tg32 and Tg32*^Ifnlr1-/-^* mice. Data shown as mean ± standard deviation with individual mice represented as points; statistical significance was determined by two-way ANOVA with multiple comparisons (**** p < 0.0001). High-magnification images in (K) and (L) are pseudocolored white and brightness adjusted to enhance visibility. Scale bars: 2.5 mm (E), 1 mm and 50 µm (H), 100 µm (K, L), 50 µm (M).

We next determined whether ventricular enlargement reflected persistent damage to the ChP barrier. Surprisingly, histological examination and IHC staining for the tight junction protein ZO-1 revealed that, during the recovery phase, the ChP epithelium of both infected Tg32 and Tg32*^Ifnlr1^*^⁻/⁻^ mice appeared largely intact and comparable to uninfected controls (**Supplemental Figure 7A-B**). These observations suggested that the ventricular phenotype was not driven by sustained barrier disruption at the ChP. Instead, examination of H&E-stained sections revealed a striking and selective abnormality in infected Tg32 mice. In these animals but not in infected Tg32*^Ifnlr1^*^⁻/⁻^ mice or uninfected controls, we observed focal loss of ependymal cells lining the ventricular walls (**Figure 7H, Supplemental Figure 7C**). Because ependymal cell loss is a well-established cause of hydrocephalus ^40^, we quantified this defect using IHC for the ependymal marker *SOX2*. Infected Tg32 mice with ventricular enlargement exhibited a profound reduction in *SOX2*-positive ependymal cells compared with uninfected controls, whereas Tg32*^Ifnlr1^*^⁻/⁻^ mice showed no significant loss (**Figure 7H-I**, **Supplemental Figure 7C**).

Several neurotropic viruses can directly infect ependymal cells and thereby induce their depletion ^41-43^. To determine whether this mechanism contributed to ependymal loss in our model, we performed RNAscope analysis for echovirus RNA together with the ependymal marker *Foxj1*. At two days post infection, viral RNA within ependymal cells was minimal in both Tg32 and Tg32*^Ifnlr1^*^⁻/⁻^ mice, indicating that direct infection was unlikely to account for the selective ependymal cell loss observed in Tg32 animals (**Figure 7J-K**). Because ependymal cells are highly sensitive to inflammatory cues ^41^, we next examined whether IFN signaling and subsequent inflammation might underlie this vulnerability. RNAscope analysis for the ISG *Ifit1* revealed more robust induction in ependymal cells of Tg32 mice compared to Tg32*^Ifnlr1-/-^*mice during acute infection (**Figure 7L**), indicating heightened IFN responsiveness in these cells. In addition to increased *Ifit1* expression, we also observed, through a combination of histological evaluation and RNAscope staining, a pronounced thickening of the ependyma in Tg32 mice that was not seen in Tg32*^Ifnlr1-/-^*mice or controls (**Figure 7M-N, Supplemental Figure 8A**). Ependymal thickening is a hallmark sign of gliosis, a common inflammatory process that occurs in ependymal cells and which is associated with cell death ^41,44^. Finally, we stained for the astrocyte marker, Gfap, to look for astrocytic processes extending into the ependymal cell layer, another sign of gliosis ^45^. We observed an increase in Gfap expression in the ependymas of both Tg32 and Tg32*^Ifnlr1-/-^* mice compared to uninfected controls (**Supplemental Figure 8B**). Together with the selective loss of ependymal cells observed in mice competent for type III IFN signaling, these data support a model in which sustained or excessive type III IFN responses indirectly promote ependymal cell attrition through inflammatory processes, leading to ventricular enlargement following viral clearance.

## Discussion

The ChP forms the blood-CSF barrier, a specialized epithelial interface that protects the brain while coordinating immune surveillance of the CNS. Despite its central role in controlling access to the CSF, the innate immune programs that operate at the ChP during viral infection have remained poorly defined. Here, by integrating human ChP organoids with a genetically tractable mouse model of echovirus infection, we uncover a previously unappreciated division of labor between type I and type III IFNs at this barrier and identify type III IFN signaling as a key driver of epithelial injury and long-term neuropathological sequelae following infection.

A central finding of this study is that the ChP epithelium mounts a robust type III IFN response to viral infection and is selectively equipped to respond to this pathway. Although this had not been previously demonstrated, it is consistent with the broader paradigm that epithelial barriers preferentially deploy type III IFNs as frontline antiviral defenses ^23,46-48^. In this regard, the ChP shares fundamental features with other epithelial and mucosal interfaces, extending the concept of type III IFN-mediated barrier immunity into the CNS. Our *in vivo* genetic dissection, however, reveals a striking divergence between IFN pathways at the ChP. Whereas type I IFNs are indispensable for controlling viral replication, limiting dissemination, and preventing mortality, type III IFNs play a fundamentally different role. Rather than conferring protection under conditions of high viral burden, sustained type III IFN signaling exacerbates epithelial injury and disrupts tight junction architecture. These findings reinforce the emerging view that type III IFNs are highly context dependent, with effects that can shift from protective to pathogenic depending on tissue state and inflammatory milieu. Notably, this behavior contrasts sharply with that described at the BBB, where type III IFNs strengthen endothelial tight junctions and limit viral neuroinvasion without directly suppressing viral replication ^26^. At the ChP epithelium, by contrast, type III IFN signaling disrupts junctional organization and impairs epithelial repair. This divergence represents an important area of future work to determine how these negative effects of type III IFN signaling may impact the permeability of the blood-CSF barrier. Together, these observations underscore that type III IFNs do not exert uniform barrier-protective effects across the brain, but instead engage distinct, tissue-specific programs with fundamentally different consequences for barrier resilience and long-term recovery.

This duality mirrors observations in peripheral tissues, where type III IFNs can restrict viral replication while simultaneously impairing epithelial repair in the lung and intestine during acute infection and chronic inflammation ^25,38,39^. Our findings extend this paradigm to the brain, identifying the ChP as a previously unrecognized site at which type III IFNs impose a barrier-disruptive state. Consistent with this model, immunohistochemical staining for markers of cell death revealed increased levels of apoptosis in the ChP of mice capable of responding to type III IFNs, highlighting their ability to induce epithelial injury. In addition, transcriptomic analysis of IFN-λ–treated ChP organoids revealed coordinated repression of gene programs governing epithelial differentiation, extracellular matrix organization, and cell–cell adhesion. Together, these data suggest that at the blood-CSF interface, type III interferons enforce a trade-off between sustained antiviral readiness and epithelial resilience.

An important implication of this work is that type III IFNs may shape CNS pathology not only through direct effects on the ChP epithelium, but also by acting on neighboring barrier-associated and immune cell populations. In addition to epithelial cells, we found that type III IFN receptors are expressed by resident ChP macrophages. These Kolmer cells are known to regulate immune cell trafficking and contribute to tissue repair following injury ^12^. While we did not observe differences in the number of macrophages in the ChP across mice during acute infection, it is plausible that type III IFN signaling skews macrophage function toward a pro-inflammatory state that is poorly suited to the barrier restoration processes necessary post infection. Defining how IFN programs influence macrophage behavior and epithelial-immune crosstalk at the ChP will be an important area for future investigation. In addition, lineage-specific knockouts for both epithelial cells and immune cells will also be a critical next step in determining how IFN signaling in these populations shapes both tissue damage and barrier integrity at the choroid plexus.

In this study, we establish a direct link between acute type III IFN responses at the ChP and long-term neurological sequelae following viral infection. Aseptic meningitis in early life is often clinically self-limited, yet a substantial proportion of survivors go on to develop persistent neurodevelopmental and cognitive impairments ^5^. The biological mechanisms that connect transient infection to delayed structural and functional abnormalities of the brain have remained poorly understood. Here, we provide evidence that type III IFN signaling, rather than direct viral cytopathology, plays a central role in driving chronic pathology of the ventricular system. In mice competent for type III IFN signaling, echovirus infection resulted in marked motor deficits, ependymal cell loss, and ventriculomegaly after viral clearance, whereas animals lacking the type III IFN receptor were protected from these sequelae. Ependymal cells are essential for CSF flow and ventricular homeostasis, and their loss is a well-established trigger of hydrocephalus ^41,49^. Although we detected little direct infection of ependymal cells, these cells exhibited robust induction of ISGs during acute infection, indicating heightened sensitivity to the inflammatory milieu generated at the blood-CSF interface. In concordance with this, we observed signs of inflammation in the ependymal cells of infected Tg32 mice that were not observed in Tg32*^Ifnlr1-/-^* mice including thickening of the cell layer as well as increased *Gfap* expression. These observations support a model in which sustained or excessive IFN exposure, originating from the ChP, perturbs ependymal cell function and viability, leading to long-term disruption of CSF dynamics. Even modest impairment of ependymal integrity can alter ventricular flow, promote ventricular enlargement, and trigger secondary neuroinflammatory processes, providing a mechanistic bridge between acute viral infection and delayed neurological disease.

Together, this study establishes the ChP as a dynamic immune barrier whose antiviral defenses are governed by a finely tuned balance between IFN pathways. We demonstrate that type I IFNs provide indispensable protection against viral replication, dissemination, and lethality, whereas type III IFNs impose a paradoxical cost by compromising epithelial integrity and predisposing the ventricular system to long-term injury. By developing complementary human and mouse models of ChP-targeted viral infection, we provide a framework for interrogating how immune responses at the blood-CSF interface shape both acute disease outcomes and chronic neurological health. More broadly, our findings redefine the role of type III IFN in the CNS, revealing that antiviral programs at barrier tissues can carry enduring structural consequences long after viral clearance.

## Figure legends

**Supplemental Figure 1. Additional characterization of echovirus tropism in human cerebral organoids.** All cerebral organoids used were 70±5 days old. **(A)** UMAPs of scRNA-seq from three mock-infected cerebral organoids collected at four days post infection split by sample. **(B)** UMAPs of scRNA-seq from three E5-infected cerebral organoids collected at four days post infection split by sample. Cell types are annotated as follows: callosal projection neurons (CPNs), corticofugal projection neurons (CFuPNs), inhibitory interneurons (Inh_INs), excitatory neurons of unknown identity (Exc_Ukn), radial glial cells (RGCs), ribosome-high cells (Ribo), choroid plexus epithelium (ChP), intermediate progenitor cells (IPCs), and retinal epithelial cells (Ret_epi). **(C)** Representative IF images from a cerebral organoid at five days post infection for DAPI (gray), vRNA (green), and *SOX2* (red). Scale bar is 50 µm. **(D)** Quantification of cells positive for both *SOX2* and vRNA from cerebral organoids across the five-day time course. Data shown as mean ± standard deviation with individual *SOX2*-positive regions within the organoids represented as datapoints.

**Supplemental Figure 2. Echovirus infection shows high specificity for the choroid plexus epithelium in mice and does not show differences in susceptibility across ventricle locations.** Tg32 mice were intracranially inoculated with 200 PFU of E5 at postnatal day three. **(A)** Representative image of a tile scanned whole brain from a Tg32 mouse at two days post infection stained via RNAscope for DAPI (gray), vRNA (green), and *Map2* (red). Yellow dashed boxes indicate regions shown at higher magnification to the right, stained for *Map2* (1), *Gfap* (2), and *Sox2* (3). **(B)** Quantification of cells positive for both vRNA and cell marker genes (*Ttr*, *Sox2*, *Map2*, and *Gfap*). Data shown as mean ± standard deviation with individual mice represented by data points. **(C)** Representative image of a tile scanned whole brain section from a Tg32 mouse at two days post infection stained via RNAscope for viral RNA (green) and DAPI (gray). Numbered dashed yellow boxes indicate zoomed regions to the right. (1) is the lateral ventricle and (2) is the fourth ventricle. **(D)** Quantification of vRNA-positive cells within the choroid plexus of Tg32 mice in both the lateral ventricles and 4^th^ ventricles. Data shown as mean ± standard deviation with individual mice represented as datapoints. No significant differences were detected by unpaired t test. Dashed yellow box indicates zoomed region to the right. Split channels have been pseudocolored white and brightness has been adjusted to enhance visibility (A and B). Scale bars are 1mm and 100um **(A-C)**.

**Supplemental Figure 3. Characterization of inflammatory and antiviral responses in human choroid plexus organoids.** All choroid plexus organoids used were 55±5 days old. Supernatants were collected at 1-, 3-, and 5-days post infection from ChP organoids infected with 200 PFU E5. Bar plots showing the induction (pg/mL) of **(A)** the matrix metalloproteases MMP-1, MMP-2, and MMP-3, **(B)** the chemokines CXCL10, CCL7, and CCL8, and **(C)** the cytokines IL-6, IL-8, and IL-34. Data are shown as mean ± standard deviation with individual organoids shown as data points. **(D)** ChP organoids were infected with 200 PFU of E5 overnight and then treated with either 0, 1,000 ng/mL recombinant human IFN-λ3, or 1,000 IU/mL of recombinant human IFNβ. Bar plot showing E5 RNA levels measured by quantitative PCR in ChP organoids following IFN treatment; data are shown as mean ± standard deviation with individual organoids represented as points. **(E)** ChP organoids were treated with either 0 or 1,000 IU/mL of recombinant human IFNβ. Bar plot showing induction of two canonical ISGs, *RSAD2* and *IFI44L*, in IFNβ treated ChP organoids compared to untreated controls. Data shown as mean ± standard deviation with individual organoids shown as data points.

**Supplemental Figure 4. Viral titers are consistent across sex.** All four genotypes (Tg32, Tg32*^Ifnlr1-/-^*, Tg32*^Ifnar1-/-^*, and Tg32*^Ifnar1-/-Ifnlr1-/-^*) were intracranially inoculated with 200 PFU of E5 at postnatal day three. **A)** Viral titers in the brain, blood, and liver across all four genotypes (Tg32, Tg32*^Ifnlr1-/-^*, Tg32*^Ifnar1-/-^*, and Tg32*^Ifnar1-/-Ifnlr1-/-^*) split by sex at 2-days post infection as determined by TCID50 assay. There was no significant difference between sex across all genotypes as determined by two-way repeated measure ANOVA. Titers are shown as log_10_TCID50 per milliliter with the limit of detection shown as a dotted line. Data are shown as mean ± standard deviation with individual animals shown as data points.

**Supplemental Figure 5. Additional histopathological characterization of the choroid plexus during infection.** (**A-E**) Uninfected Tg32 mice were intracranially inoculated with saline at postnatal day 3 and brains were harvested at two days post inoculation. **(A)** Representative images of H&E staining showing the choroid plexus of an uninfected Tg32 mouse. Dashed box shows the zoomed region to the right. **(B-E)** Representative images of IHC staining for Ki67 (**B**), CC3 (**C**), ZO-1 (**D**), and F4/80 (**E**) in the choroid plexus of uninfected Tg32 mice. **(F)** Quantification of IHC staining for F4/80 across all four genotypes of mice at two days post infection. Data shown as mean ± standard deviation with individual mice represented by datapoints; no significant differences were determined by two-way ANOVA with multiple comparisons. Scale bars are 500 μm and 100 μm **(A)** and 50 μm **(B-F)**.

**Supplemental Figure 6. Tg32 mice exhibit more severe growth deficits than Tg32*^Ifnlr1-/-^* mice. (A-C)** Tg32 and Tg32*^Ifnlr1-/-^*mice were infected with 1,000 PFU of E5 at postnatal day 3 and monitored for ∼17 days. **A)** Survival curves from mock and infected Tg32 and Tg32*^Ifnlr1-/-^* mice. **B)** Weight gain curves from mock and infected Tg32 and Tg32*^Ifnlr1-/-^* mice. **C)** Gross images of mock and infected Tg32 and Tg32*^Ifnlr1-/-^*mice at the time of collection showing size and anatomy differences.

**Supplemental Figure 7. The choroid plexus epithelium does not show signs of pathology following viral clearance. (A-B)** Tg32 and Tg32*^Ifnlr1-/-^* mice were infected with 1,000 PFU of E5 at postnatal day 3 and harvested at ∼17 days post infection. **A)** Representative images of H&E staining showing the choroid plexus from mock and infected Tg32 and Tg32*^Ifnlr1-/-^* mice. Dotted black box shows the zoomed region below. **B)** IHC-staining for ZO-1 from mock and infected Tg32 and Tg32*^Ifnlr1-/-^* mice. **(C)** Uninfected control Tg32 and Tg32*^Ifnlr1-/-^* mice were injected with saline at postnatal day 3 and monitored over time. Representative images of H&E-stained sections and serial sections IHC-stained for SOX2 from mock Tg32 and Tg32*^Ifnlr1-/-^* mice at 17 days post injection. Scale bars are 500 μm, 100 μm **(A)**, 50 μm **(B)**, and 1 mm and 50 μm for **(C).**

**Supplemental Figure 8. Tg32 and Tg32*^Ifnlr1-/-^* mice show signs of ependymal cell inflammation during acute infection. (A-B)** Tg32 and Tg32*^Ifnlr1-/-^*mice were infected with 1,000 PFU of E5 at postnatal day 3 and harvested at two days post infection. **(A)** Representative H&E images of ependymal cells of uninfected and infected Tg32 and Tg32*^Ifnlr1-/-^* mice at two days post infection. **(B)** Representative RNAscope stained images of ependymal cells from uninfected and infected Tg32 and Tg32*^Ifnlr1-/-^* mice stained for DAPI (gray) and *Gfap* (green). Values in the lower left corner represent total *Gfap* signal intensity. Scale bars: 50 μm **(A-B)**.

## Materials and methods

### Cell lines and viruses

HeLa cells (clone 7B) were provided by Jeffrey Bergelson, Children’s Hospital of Philadelphia (Philadelphia, PA), and cultured in minimal essential medium (MEM) supplemented with 5% fetal bovine serum (FBS), nonessential amino acids, and penicillin-streptomycin. Experiments were performed with E5 (Noyce strain), which was obtained from the ATCC. Virus was propagated in HeLa 7B cells and purified by ultracentrifugation over a 30% sucrose cushion, as described previously ^50^. The purity of viral stocks was confirmed by deep sequencing.

### Animals

All animal experiments were approved by the Duke University Animal Care and Use Committees, and all methods were performed in accordance with the relevant guidelines and regulations. All four genotypes of mice including Tg32, Tg32*^Ifnlr1-/-^*, Tg32*^Ifnar1-/-^*, and Tg32*^Ifnar1-/-Ifnlr1-/-^* had been previously described ^24,34^. All animals used in this study were genotyped by Transnetyx, and genotyping assay results are available upon request.

### Human cerebral organoid and choroid plexus organoid culture

Cerebral and choroid plexus organoids were grown from either human embryonic stem cells (WA09) or human induced pluripotent stem cells (SCTi005) as previously described ^27^. Briefly, embryoid bodies (EBs) were generated by seeding 2,000 stem cells per well in a 96-well U-bottom ultra-low attachment plate in hESC media containing 4 ng/mL of FGF2 and 50 μM of ROCK inhibitor for four days. At day 5 the media was refreshed with hESC media without FGF2 and ROCK inhibitor. At day 7 each EB was moved to its own well of a 24-well ultra-low attachment plate containing 500 μL of NI media. At day 10 EBs were embedded in 30 μL of Matrigel using dimpled parafilm and then released into suspension in a 60 mm dish containing 5mL of cerebral organoid differentiation media without vitamin A. At day 14 the dishes containing the organoids were moved to an orbital shaker (65 r.p.m.) and the media was replaced with cerebral organoid differentiation media with vitamin A. Cerebral organoids were maintained in these conditions for maturation and growth with media changes every 2-3 days. For choroid plexus organoids at days 10-17, 20 ng/mL BMP4 and 3 μM CHIR were added to the cerebral organoid differentiation media, as described ^13^.

### Choroid plexus organoid area measurement

Choroid plexus organoids were infected with 200 PFU of E5 and total area measurements were taken each day following infection. Area measurements were taken using a Keyence BZ-X810 microscope. Briefly, images of the entire organoid were taken and then the total area measurements were made by drawing a region of interest around the entire organoid. Then, the automated measure feature of the Keyence image analysis software was used to calculate the total area of the organoid.

### IF staining

Organoids were fixed in 4% paraformaldehyde (PFA) overnight at 4°C and then moved to a 30% sucrose buffer. The organoids were then embedded in O.C.T. and flash frozen. Blocks were stored at -80°C. Sections were cut at a thickness of 10 μm and frozen slides were stored at -20°C. Prior to staining, sections were incubated with 4% PFA for 10 minutes and then washed with 1× DPBS. Sections were then incubated with a permeability/blocking buffer (10% normal goat serum and 0.5% Triton X-100 in 1× PBS) for 45 minutes at RT. Sections were then incubated with primary antibody diluted in primary antibody diluent (10% normal goat serum and 0.1% Triton X-100 in 1× DPBS) overnight at 4°C. A detailed list of the antibodies used is included in the supplementary materials **(Supplemental table 5)**. Sections were incubated with secondary antibodies for 2 hours at RT. Images were taken using an Olympus Fluoview FV3000 confocal microscope or a Keyence BZ-X810 microscope and images were prepared and/or brightness contrast adjusted using Fiji.

### Single cell dissociation

Cerebral organoids at four days post infection were washed extensively with 1× DPBS before being incubated with 1 mL of TrypLE Express (ThermoFisher, cat. #12605010) on the orbital shaker for 15 min in the 37°C incubator. Cerebral organoids were then manually disrupted using cut pipet tips with progressively smaller openings. TrypLE Express was then inactivated by adding 1 mL of cerebral organoid differentiation media with vitamin A. The cell suspension was then passed through a 30 μm cell strainer to remove any large clumps and debris. The filtered cell suspension was then spun down at 400 × g for 5min, and the cells were resuspended in 1× DPBS. Dead cells were then removed using the Dead cell removal kit (Miltenyi Biotec, cat #: 130-090-101). Finally, the resulting cell suspension was spun down at 400xg for 5min and then resuspended in 1% bovine serum albumin (BSA) in 1× DPBS. The samples were then placed on ice and taken to the Duke sequencing core for scRNAseq sample processing.

### Single cell analysis

Libraries were prepared from 10k total cells using the methods described in the 10X Genomics Single Cell 3’ Reagent Kit Protocol (v2 chemistry, Manual Part #CG00052). Sequencing runs were performed on an Illumina NovaSeq XPlus (Illumina, San Diego) using an S2 flow cell (Illumina, San Diego), which allowed for ∼62k reads/cell. FASTQ files were uploaded to the 10X Cloud Analysis website (https://www.10xgenomics.com/products/cloud-analysis) and analyzed using Cell Ranger v7.1.0 using a custom reference genome including the human genome (GRCh38) and the echovirus-5 genome. Mapped files were downloaded through command line and filtered feature matrices utilized for secondary analyses in R (v4.3.1) using a Seurat-based pipeline. Cells were retained if they contained 200–8,000 detected genes, fewer than 30,000 total counts, and less than 7.5% mitochondrial transcripts. Following QC, individual filtered matrices were combined for both the infected and uninfected samples. The gene *MALAT1* was removed from both combined files as a part of QC. SCTransform normalization was applied to each dataset, regressing out mitochondrial content, ribosomal content, and gene/UMI counts. Analysis code is available on the CoyneLab GitHub repository (https://github.com/CoyneLabDuke). All raw files have been deposited on SRA under the following BioProjectID: PRJNA1402435.

### CSF collection

Choroid plexus organoids were washed extensively with 1× DPBS to minimize residual culture medium. CSF-like fluid was collected using an insulin syringe by gently aspirating fluid from the center of organoid cysts. Samples were stored at −20 °C until analysis.

### Luminex

Luminex profiling was performed on culture supernatants and CSF-like fluid collected from choroid plexus organoids. A human inflammation panel kit (171AL001M, Bio-Rad) and a human chemokine panel kit (171AK99MR2, Bio-Rad) were used according to the manufacturer’s instructions. Assays were run on a Bio-Plex 200 system (Bio-Rad), and data were acquired and analyzed using Bio-Plex Manager software (Bio-Rad). Heat maps were generated using fold changes in average analyte concentration (pg/mL) at each time point relative to uninfected controls and were plotted using GraphPad Prism.

### IFN and poly(I:C) treatment, qPCR, and Bulk RNA Sequencing of Choroid Plexus Organoids

For poly(I:C) stimulation experiments, ChP organoids (∼55 days in culture) were treated overnight with 10 μg/mL poly(I:C) (Invivogen, Cat #tlrl-pic) diluted in cerebral organoid differentiation media containing vitamin A. The following day, organoids were harvested for RNA extraction.

For pretreatment experiments, ChP organoids were treated overnight with 0, 1,000 IU/mL recombinant human IFNβ (rhIFNβ; R&D Systems, cat. No. 8499-IF-050/CF), or 1,000 ng/mL recombinant human IFN-λ3 (rhIFN-λ3; R&D Systems, cat. no. 8417-IL-025/CF) diluted in cerebral organoid differentiation medium containing vitamin A. The following day, organoids were inoculated with 200 PFU of E5 and incubated for 2 h at 37 °C. After incubation, the inoculum was removed and replaced with fresh differentiation medium containing the corresponding IFN, and organoids were returned to the incubator. Culture medium was replaced daily with fresh IFN-containing medium. Organoids were harvested at three days post infection for RNA extraction.

For post-infection treatments, ChP organoids were inoculated with 200 PFU of E5 and incubated for 2 h at 37 °C. After incubation, the inoculum was removed and replaced with fresh differentiation medium overnight. The following day, media was replaced with fresh differentiation media containing either 0, 1,000 IU/mL rhIFNβ, or 1,000 ng/mL rhIFN-λ3. Culture medium was replaced daily with fresh IFN-containing medium. Organoids were harvested at three days post infection for RNA extraction.

Total RNA was isolated from choroid plexus organoids using the GenElute Mammalian Total RNA Miniprep Kit (MilliporeSigma, RTN350) according to the manufacturer’s instructions. RNA was reverse transcribed into cDNA using the iScript cDNA Synthesis Kit (Bio-Rad, 1708891). Quantitative PCR (qPCR) was performed using iTaq Universal SYBR Green Supermix (Bio-Rad, 1725121) on a Bio-Rad Opus real-time PCR system. Primer pairs used were human IL28A (MilliporeSigma; forward 04250482-001, reverse 04250483-001), human IL29 (MilliporeSigma; forward 05010044-001, reverse 05010045-001), human IFNB1 (Qiagen; QT00203763), EVB (IDT; forward 564892784, reverse 564892785), human RSAD2 (MilliporeSigma; forward 01233280-001, reverse 01233281-001), human IFI44L (MilliporeSigma; forward 02606353-001, reverse 02606354-001), and human ACTB (MilliporeSigma; forward SY190735585-067, reverse SY190735585-068). Gene expression was normalized to ACTB using the ΔCt method. Relative expression was calculated using the ΔΔCt method, with untreated control samples used as the reference condition, and fold changes in expression were determined as 2^−ΔΔCt.

For bulk RNA sequencing, library preparation was performed by the Duke Center for Computational Biology and Genomics using a stranded mRNA KAPA HyperPrep kit. Sequencing was carried out on an Illumina NovaSeq X Plus platform using 100-bp paired-end reads, targeting a total of approximately 10 billion reads across all samples. Reads were aligned to the human genome using Qiagen CLC Genomics Workbench (v20). Differential gene expression analysis was performed using DESeq2 in R. Raw count data were normalized and analyzed to identify differentially expressed genes between conditions ^51^. Genes with an adjusted p value < 0.05 and an absolute log₂ fold change ≥ 1 were considered significantly differentially expressed.

### Mouse infections

Neonatal mice at postnatal day three were inoculated intracranially (i.c.) with either 200 or 1,000 PFU of E5 virus. Mice were anesthetized by hypothermia on ice until movement ceased and then placed on a chilled aluminum surface to maintain anesthesia throughout the injection procedure. Animals were injected with 2 μL of either sterile 1× phosphate-buffered saline (DPBS; mock-infected) or virus diluted in 1× DPBS using a 10 μL Hamilton syringe fitted with a 32-gauge needle. Injections were performed into the right cerebral hemisphere, approximately 2 mm lateral to the intersection of the sagittal and lambdoid sutures and at a depth of approximately 2 mm. Following injection, mice were placed on a warming pad and monitored until full recovery. Animal health and survival were assessed every 24 h for the duration of the experiment.

Mice were euthanized at the indicated time points, and organs were harvested into tissue homogenization tubes containing 0.5 mL of HeLa growth medium and stored at −80 °C. For viral titration, tissues were thawed and homogenized using a Bead Ruptor Elite bead mill homogenizer (Omni International) at 4 m/s for one cycle of 20 s for mice younger than 14 days of age or one cycle of 1 min for mice older than 14 days. Homogenates were clarified by centrifugation at 10,000 × g for 5 min to pellet debris. Viral titers in tissue homogenates were determined by 50% tissue culture infective dose (TCID50) assays on HeLa cells and quantified following crystal violet staining.

For surface righting reflex tests, pups were placed on their backs on a flat surface covered with a paper towel. Pups were released and timed with a stopwatch until they were able to flip over with all four paws flat on the surface. The measurement was repeated three times per pup with a 30 second rest period between each measurement.

### Histology and immunohistochemistry

All histological and immunohistochemical staining was performed by HistoWiz, Inc. using a Leica Bond RX automated stainer (Leica Microsystems) with a fully automated workflow. Mouse brains and organoids were fixed in 10% neutral-buffered formalin for 24 h, transferred to 70% ethanol, and shipped to HistoWiz for paraffin embedding and sectioning. Tissue sections were stained with hematoxylin and eosin (H&E) or processed for immunohistochemistry using antibodies against AQP1 (Abcam, ab15080), MAP2 (Abcam, ab183830), SOX2 (Cell Signaling Technology, 14962), Ki67 (Cell Signaling Technology, 34330), F4/80 (14-4801-82), CC3 (Cell Signaling Technology, 9661), and ZO-1 (Abcam, ab221547) (see **Supplemental Table 5**).

### Staining quantification and intensity measurements

Quantification of infected cells within the choroid plexus was performed using QuPath software. For each mouse genotype, a region of interest (ROI) was drawn around the choroid plexus located in the right 4^th^ ventricle of the brain. This region of choroid plexus was selected due to its consistency across brain sections. The cell detection function was used to determine the total number of cells within the ROI, and then the positive cell detection feature was used to determine the number of virus-positive cells within the ROI for each animal. Quantification of ependymal cell layer thickness was performed using QuPath. Briefly, an ROI was drawn around the area staining positive for *Foxj1* and the cell detection feature was used to count the number of cells in the region. The number of cells spanning the thickest area of the ROI was recorded as a measure of ependymal layer thickness. This measurement was repeated for three separate areas for each mouse and averaged together. Quantification of staining intensity was performed using FIJI. Briefly, images were first thresholded to exclude background signal. A region of interest encompassing the target area was then manually defined, and integrated density was measured for the signal within the selected region.

### Morphometric and histological quantification

Post-mortem images of mice were acquired at the conclusion of each experiment with a ruler included for scale. Body length was measured from the tip of the nose to the rump, and head length was measured from the snout to the posterior aspect of the skull. Ventricular area was assessed from sagittal brain sections collected at approximately 17 days post infection. Measurements were performed using QuPath software. For each brain, the largest ventricle was identified, and a region of interest (ROI) was manually traced along the ventricular boundary. Ventricular area was calculated as the area within the ROI. Ventricles were classified as enlarged if the measured area exceeded twofold the mean ventricular area of mock-infected controls.

Quantification of IHC staining for Ki67, CC3, and F4/80 was performed using QuPath. For all measurements, an ROI was drawn around the choroid plexus in each ventricle of the brain, and the positive cell detection feature was used to determine the number of positive cells within the ROIs. The percentage of positive cells detected in each ventricle were then averaged together to get an average percent positive cells for each mouse. Quantification of ZO-1 localization was performed using QuPath. Two ROIs were drawn for each measurement: one along the apical surface of the epithelial cells and one directly underneath it, passing through the epithelial cell body. The total staining intensity for ZO-1 was measured in both ROIs using the classifier function and creating a pixel thresholder. The measurements were then used to create a ZO-1 localization ratio to evaluate disruption of tight junctions. ZO-1 staining is sharply delineated along epithelial cell borders in healthy tissue and becomes more diffuse and cytoplasmic when tight junctions are disrupted, therefore tissue with a lower ZO-1 localization ratio indicates higher levels of disruption. Quantification of SOX2-positive ependymal cells was performed using QuPath. An ROI was drawn along the ventricular wall, restricted to the ependymal cell layer. The positive cell detection function was used to determine the number of SOX2-positive cells within the ROI for each animal. Quantification of virus-infected ependymal cells was performed using FIJI. An ROI was drawn along the ventricular wall to encompass Foxj1-positive ependymal cells. Total nuclei within the ROI were counted to determine the total number of ependymal cells, and cells positive for viral RNA were enumerated within the same region. The percentage of infected ependymal cells was calculated as the number of virus-positive cells divided by the total number of ependymal cells within the ROI.

### RNAscope

RNAscope was performed as previously described ^52^. Formalin-fixed, paraffin-embedded (FFPE) tissue sections were baked at 60 °C for 1 h, deparaffinized by sequential incubation in xylene and 100% ethanol, and treated with hydrogen peroxide for 10 min at room temperature. Slides were then subjected to target retrieval in RNAscope target retrieval buffer using a pressure cooker (Instant Pot) at >99 °C for 15 min, followed by incubation in 100% ethanol for 3 min at room temperature. Protease treatment was performed at 40 °C for 30 min. During protease incubation, RNAscope probes were prewarmed at 40 °C for 10 min and allowed to equilibrate to room temperature for 20 min. Diluted probe solution was then applied to each slide, and hybridization was carried out for 2 h at 40 °C. Slides were subsequently stored overnight in 5× SSC buffer. On day 2, signal amplification was performed by sequential incubation with AMP1, AMP2, and AMP3 reagents at 40 °C for 30 min, 30 min, and 15 min, respectively, with washes between each step. Slides were then incubated with channel-specific HRP reagents (C1-HRP) for 15 min at 40 °C, followed by fluorophore development using TSA Vivid Fluorophore 520 (1:1500 dilution in TSA buffer) for 30 min at 40 °C. HRP activity was quenched by incubation with HRP blocker for 15 min at 40 °C. This amplification and detection process was repeated sequentially for each additional channel using distinct fluorophores. Following signal development, slides were treated with an autofluorescence quenching reagent (Vector Laboratories, SP-8400) for 5 min at room temperature, counterstained with DAPI for 30 s, and mounted with VECTASHIELD antifade mounting medium (Vector Laboratories, H-1200-01). Slides were imaged using either an Olympus Fluoview FV3000 confocal microscope or a Keyence BZ-X810 microscope. Image processing and analysis were performed using FIJI. A detailed RNAscope protocol is available from Advanced Cell Diagnostics (ACDbio). Probe information is provided in **Supplemental Table 5**.

## Acknowledgments

We would like to acknowledge the assistance of the Molecular Genomics Core at the Duke Molecular Physiology Institute, Duke University School of Medicine, for the generation of data for the manuscript. We thank Megan Baldridge (Washington University) for providing *Ifnlr1^-/-^*mice and Sujan Shresta (La Jolla Institute for Immunology) for providing hFcRn^Tg32^-*Ifnar1^-/-^* mice. This project was supported by NIH R01-AI150151 (C.B.C). The funders had no role in study design, data collection and analysis, decision to publish, or preparation of the manuscript.

